# The association between working memory precision and the nonlinear dynamics of frontal and parieto-occipital EEG activity

**DOI:** 10.1101/2022.04.24.489284

**Authors:** Wen-Sheng Chang, Wei-Kuang Liang, Dong-Han Li, Neil G. Muggleton, Prasad Balachandran, Norden E. Huang, Chi-Hung Juan

**Affiliations:** Institute of Cognitive Neuroscience, College of Health Sciences and Technology, National Central University, Taiwan; Cognitive Intelligence and Precision Healthcare Center, National Central University, Taiwan; Institute of Cognitive Neuroscience, University College London, London, UK; Department of Psychology, Goldsmiths, University of London, London, UK; Data Analysis and Application Laboratory, The First Institute of Oceanography, Qingdao, China; Department of Psychology, Kaohsiung Medical University, Taiwan; Taiwan International Graduate Program in Interdisciplinary Neuroscience, National Cheng Kung University and Academia Sinica, Taipei, Taiwan

**Author notes:** Corresponding author contact information: Chi-Hung Juan Institute of Cognitive Neuroscience, College of Health Sciences and Technology, National Central University, Taiwan. **Institutional Review Board Statement** This study was approved by the Institutional Review Board of the National Taiwan University, Taipei, Taiwan (Ethical Approval Code is 201710EM023). Informed consent was obtained from all participants involved in the study. Abbreviation Table: AM: amplitude modulation; AMF: amplitude-modulation frequency; CF: carrier frequency; CFC: cross-frequency coupling; DQ: direct quadrature; EMD: empirical mode decomposition; FDR: false-discovery rate; FO: fast oscillation; GLM: general linear model; HHCFPC: Holo-Hilbert cross-frequency phase clustering; HHS: Holo-Hilbert spectrum; HHSA: Holo-Hilbert spectral analysis; IMF: intrinsic-mode function; PAC: phase-amplitude coupling; PLV: phase-locking value; SO: slow oscillation; WM: working memory.

**Keywords:** visual working memory, memory precision, Holo-Hilbert Spectral Analysis, amplitude modulation, cross-frequency coupling, neural oscillations

## Abstract

Electrophysiological working memory (WM) research has shown that distinct brain areas communicate through macroscopic oscillatory activities across multiple frequency bands. Such cross-frequency interactions generate nonlinear amplitude modulations (AM) in the observed signal. Traditionally, the AM of a signal is expressed as coupling strength between the signal and a pre-specified modulator at a lower frequency. Therefore, the idea of AM and coupling cannot be separately studied. This EEG study shows that the AM of parieto-occipital alpha/beta power and the coupling between frontal theta phase and parieto-occipital alpha/beta AM provide different information on WM processing. Thirty-three participants completed a color recall task with simultaneous EEG recording. The results showed that individual differences in WM precision are associated with frontal theta power enhancement and parieto-occipital alpha/beta power suppression. Furthermore, the AM of parieto-occipital alpha/beta power predicted WM precision after presenting a target-defining probe array. The phase-amplitude coupling (PAC) between frontal theta phase and parieto-occipital alpha/beta AM increased with WM load during the processing of incoming stimuli, but they did not predict the subsequent recall performance. These results indicate that the frontoparietal PAC reflects the executive control for selecting relevant WM representations, but whether the memorized information can be retrieved depends on the subsequent amplitude variation of parieto-occipital alpha/beta power. In conclusion, individuals with higher working memory precision are associated with enhanced frontal theta power and parieto-occipital alpha/beta power suppression.

## Introduction

WM enables us to store and utilize information in the absence of external stimuli for a brief period. Previous studies have associated WM performance with oscillatory activities and interareal connectivity between frontal and posterior brain regions (D’Esposito and Postle, 2015; Dipoppa et al., 2016; Eriksson et al., 2015; Gazzaley and Nobre, 2012). Specifically, alpha/beta (8-30 Hz) power suppressions in task-relevant sensory areas and frontal theta (4-8 Hz) power increment during WM retention periods are frequently reported (van Ede et al., 2017; Erickson et al., 2019; Fukuda et al., 2015, 2016; Hsieh and Ranganath, 2014). Suppression of alpha/beta power is often associated with the processing involved in the active maintenance of memory representations (van Ede, 2018; Foster et al., 2016; Fukuda et al., 2016; Griffiths et al., 2019), whereas frontal theta power reflects the controlling process during memory tasks (Hanslmayr and Staudigl, 2014; de Vries et al., 2020).

Recent advances in electrophysiological WM research have focused on interactions across different frequency components and brain areas (Daume et al., 2017; Johnson et al., 2017; Liang et al., 2021; Popov et al., 2018; Roux and Uhlhaas, 2014; de Vries et al., 2018). Measures of cross-frequency coupling (CFC) are often used to quantify these WM-related cross-frequency interactions, e.g., the phase-amplitude coupling (PAC) between fast and slow oscillations (Canolty and Knight, 2010; Tort et al., 2010). In PAC, fast and slow oscillatory components are filtered from the raw signal, then the phase of the slower component is compared to the amplitude of the faster component with synchronous measures. Typically, stronger PAC is associated with greater WM demands or better WM performance (Axmacher et al., 2010; Daume et al., 2017; Friese et al., 2013; Roux and Uhlhaas, 2014; Siegel et al., 2009).

Although frontal theta power enhancement and parieto-occipital alpha power suppression during WM maintenance have been frequently reported, their coupling is rarely discussed. Indirect evidence (e.g., Granger causality) have shown that frontal to parieto-occipital theta connectivity is transiently modulated by incoming executive demands, whereas parieto-occipital to frontal alpha/beta connectivity is sustained throughout the task procedure (Johnson et al., 2017; Popov et al., 2018). One recent dual-task study reported that frontal-midline theta power predicted the subsequent parieto-occipital alpha power lateralization, furthermore, enhanced coupling was observed between frontal-theta phase and parieto-occipital alpha amplitude in the cued region (de Vries et al., 2018). Nevertheless, the correlation between frontoparietal theta-alpha PAC and subsequent recall performance remained unexplored.

The standard PAC methods are increasingly criticized due to potential confounding factors arising from the nonstationary and nonlinear nature of biological signals (Aru et al., 2015; Cole and Voytek, 2017; Hyafil, 2015). The most fundamental issue comes from the physical meaning of PAC, which is a multiplicative process that cannot directly be measured with additive time-frequency decomposition methods such as short-time Fourier and wavelet decompositions (Huang et al., 2016; Juan et al., 2021). The same limitation holds for bispectral measures frequently used to study the interactions between various frequency components of a signal (Kovach et al., 2018; Shahbazi Avarvand et al., 2018). In these methods, the amplitude modulation (AM) of a fast wave is measured by its projection to a reference slow wave, so that PAC and AM are inseparable concepts. As a result, the rhythmic AM of a fast wave cannot be measured if the frequency-matched slow wave does not present in the observed data (Fig. 1). Such constraint limits the scope of studying the nonlinear properties of oscillatory signals.

**Figure 1.**
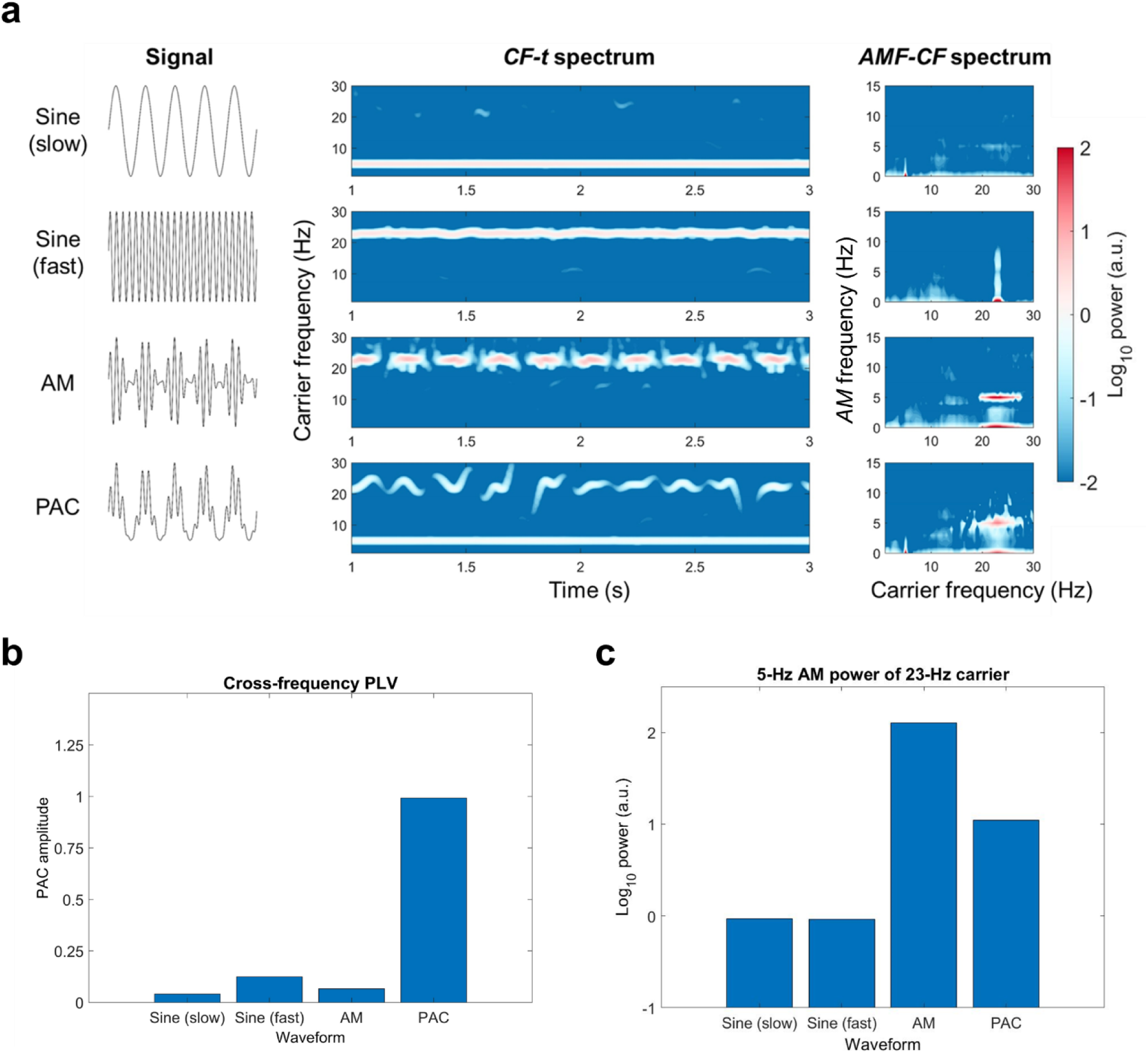
Comparison of PAC amplitude and AM power in different waveforms. (a) Simulated waveforms and their spectral representations. The waveforms are decomposed by HHSA. (*Left column*) The tested waveforms are 5-Hz sine wave, 23-Hz sine wave, 23-Hz sine wave with 5-Hz AM, and a PAC waveform which is composed of a 5-Hz sine wave and a 23-Hz sine wave with 5-Hz AM. (*Middle column*) The time-frequency representation of the simulated waveforms. The rhythmic intermittent patterns for the AM and PAC waves indicate the existence of AM for the 23-Hz carrier. (*Right column*) The *AMF-CF* representation of the simulated waveforms. The horizontal axis denotes carrier frequency, and the vertical axis denotes the AM frequency. The energy of the fast and the slow sine waves are concentrated in their corresponding carrier frequencies without AM (i.e., 0 Hz). For the AM wave, the energy is concentrated at 23-Hz carrier frequency both at the DC level and 5-Hz AM frequency. An additional 5-Hz carrier-frequency component can be observed at the DC level for the PAC wave. (b) The power at 23-Hz carrier frequency with 5-Hz AM for the four waveforms tested. (c) PAC amplitudes between 5-Hz and 23-Hz components. Only the PAC wave shows large PAC amplitude.

The main purpose of the study is to investigate how the AMs of EEG oscillation is associated with WM performance. Thirty-three participants took part in a color recall task with concurrent EEG recording (Fig. 2a). On each trial, participants were required to remember a set of color squares. After one second, a set of framed squares was presented at the same location, and the retrieval target was defined as the one on the location with a thicker frame. Participants made responses by matching the color with a color wheel that was presented after another second. WM precisions under different WM loads served as an index of WM performance (Fig. 2b, Bays et al., 2009; Zhang and Luck, 2008). The observed EEG data was decomposed with *Holo-Hilbert spectral analysis* (HHSA) that can directly measure the AM energy in nonlinear signals (Huang et al., 2016). In HHSA, the AMs of signal carriers are decomposed into different amplitude-modulating frequencies (AMF). Therefore, not only the *carrier frequency (CF)—time* profile, but also the *AMF—CF* structure can be revealed with the corresponding Holo-Hilbert spectra (HHS). We conducted separate analyses for nonlinear EEG variations following the load manipulation and how EEG variation is correlated to individual differences in WM precision after controlling for WM loads. After identifying the oscillatory correlates of WM precision, we further examined whether there are any systematic variations in frontoparietal PAC.

**Figure 2.**
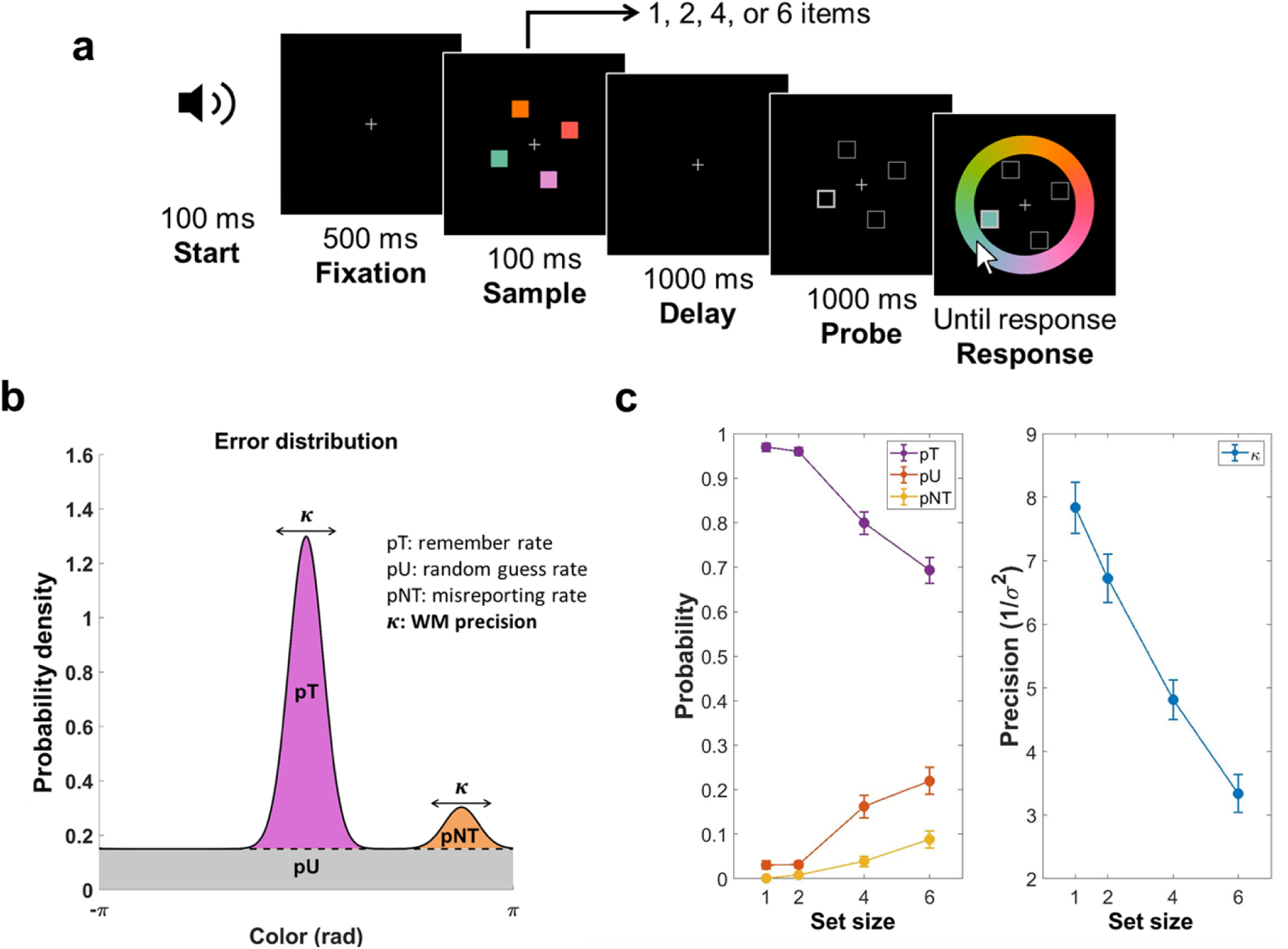
Design and behavioral results of the color recall task. (a) Participants were instructed to remember the color of each square in the sample array. After a 1000-ms delay period, a set of framed squares were presented to indicate sample locations. The target item was defined as the one with bold edges, and participants had to match the target color with a color wheel. During the response period, the color at the target location kept updating according to selected color from the mouse cursor. The set size of the sample array was manipulated randomly in four conditions (1, 2, 4, or 6 items, interleaved design). (b) A mixture model of three components modeled the response error distribution of each condition. First, the memory of item colors was stored with noise, therefore the response error distribution could be explained by a von Mises distribution with concentration number κ (reciprocal of the error variance, 1/*σ*^2^), centering at the target color location (*purple*). Since the target was not determined during the sample array, all item colors were assumed to be encoded with equal precision. Second, there was a probability of misreporting non-target colors. In this case, the variability of the response errors was explained by a set of von Mises distributions centering at the location of non-target colors with the same κ value as the remembered situation (*orange*). Third, when participants made random responses, the errors would be uniformly distributed (*gray*). Ratios of the three components could be estimated by two scalars pU (the probability of random guessing) and pNT (the probability of misreporting non-target colors), pT (the probability of correctly remembering the target color) equaled 1-pU-pNT. The results of estimated mixture-model parameters were illustrated in (c) and (d). (c) pT decreased (purple line; *F_(3,96_*_)_ = 55.52, *p* < 10^-5^; *η*^2^ = 0.6344), whereas pU (red line; *F_(3,96)_* =26.20, *p* < 10^-5^; *η*^2^ = 0.4501) and pNT (green line; *F_(3,96)_* = 12.05, *p* < 10^-5^; *η*^2^ = 0.2764) increased as set size increased. (d) WM precision was defined as the concentration number κ. κ decreased (*F_(3,96)_* = 48.54, *p* < 10^-5^; *η*^2^ = 0.6027) as set size increased.

## Results

### Comparison of AM and PAC

Here we use a simulation to address the difference between AM and PAC (Fig. 1; see also Juan et al., 2021). The simulation compares the spectral representations of an AM signal, a PAC signal, and their constituent sine waves. The spectral representations of the signals are generated with HHSA (Huang et al., 2016). In this simulation, the high-frequency component of the AM and the PAC signals are modulated in the same frequency, but only the PAC signal contains the frequency-matched slower components (Fig. 1a). However, the standard PAC method (e.g., Canolty and Knight, 2010) only extracts the AM energy in the PAC waveform, despite both the AM and PAC waveforms both contain AM energy in the same modulating frequency (Figs. 1b and 1c).

### Behavioral Results

Thirty-three participants performed a color-recall task with varying WM loads. Response errors in different loads were fitted separately by a mixture model (Fig. 2b, see Materials and Methods for details). As expected, participants performed worse as set size increased (Fig. 2c). With increasing set size, the probability for remembered response (pT) decreased (*F_(3,96)_* = 55.52, *p* < 10^-5^, η^2^ = 0.6344), whereas the probability for misreported responses (pNT; F_(3,96)_ = 12.05, *p* < 10^-5^; η^2^ = 0.2764) and guessed responses (pU; *F_(3,96)_* = 12.05, *p* < 10^-5^; η^2^ = 0.2764) increased. On top of the decreasing recall rates, WM precision (κ) also decreased as set size increased (*F_(3,96)_* = 48.54, *p* < 10^-5^, η^2^ = 0.6027). Post hoc analyses revealed significant differences of κ between set sizes 1 and 2 (*t_32_* = -2.728, *p* = 0.0103) but no significant difference for pT (*t_32_* = -1.2182, *p* = 0.2321), indicating that WM precision (κ) was more sensitive in small WM loads. Parameters κ, pT, and pU show significant differences between set sizes 2 and 4 (κ, *t_32_* = -4.7273, *p* = 4.38×10^-5^; pT, *t_32_* = -7.4262, *p* = 1.89×10^-8^; pU, *t_32_* = 5.7989, *p* = 1.94×10^-6^), as well as 4 and 6 (κ, *t_32_* = -3.889, *p* = 4.78×10^-4^; pT, *t_32_* = -4.6453, *p* = 5.55×10^-5^; pU, *t_32_* = 2.581, *p* = 0.0146) after FDR correction at .05 level. pNT differences between set sizes 2 and 4 (*t_32_* = 2.3583, *p* = 0.0246) as well as set sizes 4 and 6 (*t_32_* = 2.4185, *p* = 0.0215) were not significant after FDR correction.

### WM load modulated parieto-occipital alpha/beta power

The HHSA is based on the empirical mode decomposition (EMD) which decomposes the temporal signal into a set of intrinsic-mode functions (IMF, Huang et al., 1998). The IMFs serve as a dyadic filter bank to the data (Deering and Kaiser, 2005). Therefore, each IMF represents EEG activity in a distinct dyadic timescale (Fig. 3a). Our EEG analysis first verified whether the task used in the study replicated the load-dependent alpha power suppression in the previous literature (van Ede, 2018; Erickson et al., 2019; Fukuda et al., 2015; Proskovec et al., 2019). For each IMF, the grand-average amplitudes across all electrodes were calculated first, and the effect of load manipulation was evaluated by performing a repeated-measures ANOVA at each time point and each IMF. They were corrected with a cluster-based permutation procedure of 5000 iterations. The set-size 1 condition served as the baseline for comparison and the effects were quantified as the deviation from the baseline. Significant clusters were observed in IMFs 4-7, in which IMFs 4 (11.8-23.6 Hz) and 5 (6.4-11.8 Hz) showed negative load effects in the delay and probe periods, whereas IMFs 6 (3.1-6.4 Hz) and 7 (1.5-3.1 Hz) showed transient positive load effects following the onsets of sample and probe arrays (Fig. S1). For the IMF capturing alpha activity, two negative clusters could be observed within the latency of interest (−0.5 to 2.5 s relative to the onset of the sample array, Fig. 3b). The first cluster was observed in 0.5-1.15 s (cluster-corrected *p* = 4×10^-4^), and the second cluster was observed in 1.55-2.1 s (cluster-corrected *p* <= 2×10^-4^). A marginal positive load effect was observed following the onset of the sample array (0-0.2 s; cluster-corrected *p* = 0.057). On average, the F-statistics from 0 to 2.1 s after the onset of the sample array concentrated in parieto-occipital electrodes (Fig. 3c).

**Figure 3.**
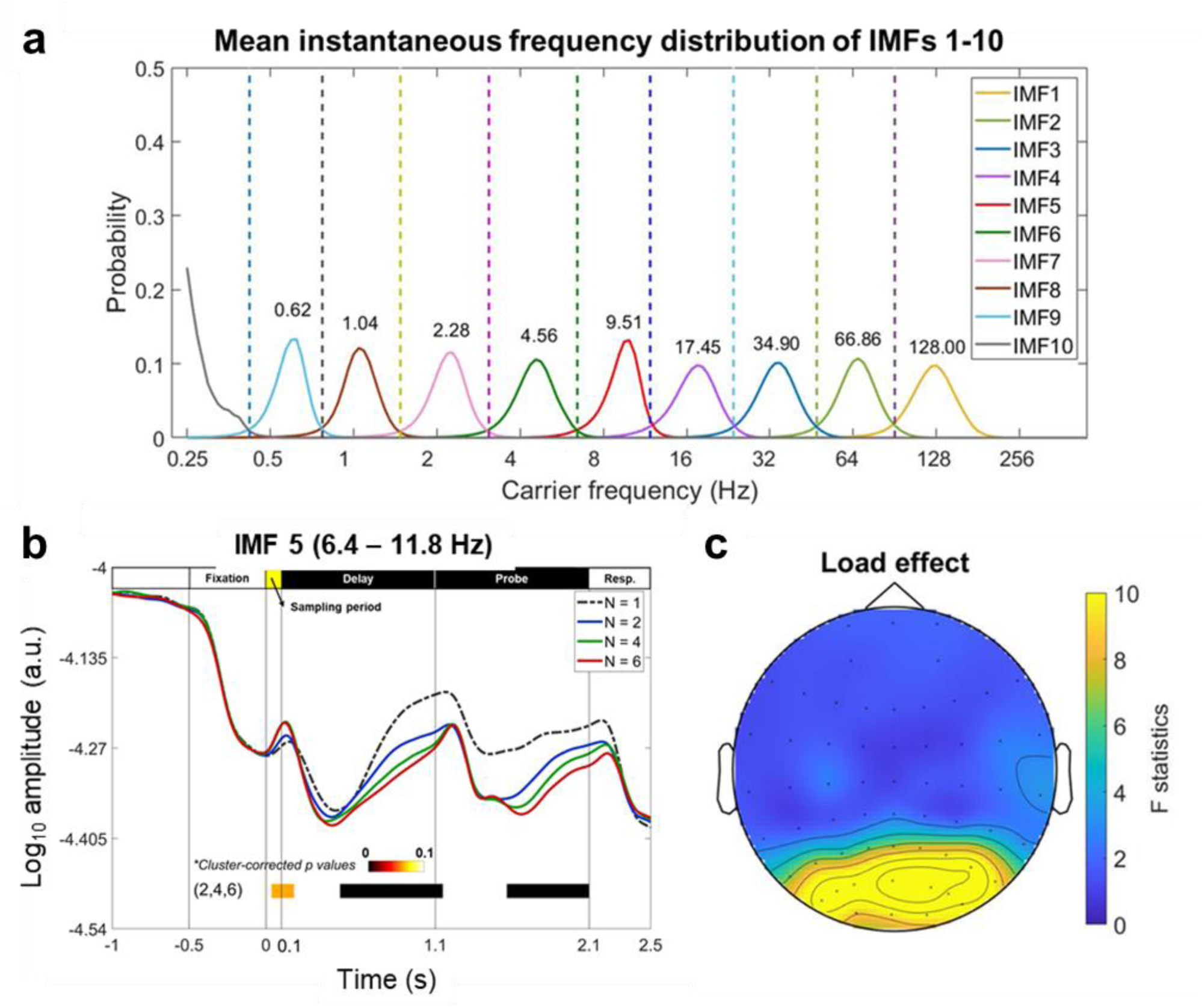
WM load modulated parieto-occipital alpha amplitude. (a) The instantaneous frequency distributions of IMFs averaged across all electrodes and all participants. Numbers denote the peak frequency of each IMF distribution. IMF 5 (6.4-11.8 Hz) fell within the timescale of the alpha-band frequency. (b) The time course of the grand average amplitude of IMF 5 across all electrodes for each set size. A repeated-measure ANOVA for set sizes 2, 4, and 6 (set-size 1 served as the baseline condition for comparison) was performed for each time point then was corrected by a cluster-based permutation procedure of 5000 iterations. Color bars indicated cluster-corrected *p* values for each cluster. Three significant clusters were observed. A positive cluster could be identified following the sample array onset (cluster-corrected *p* = 0.057), and two negative clusters could be observed both in the delay (cluster-corrected *p* = 4×10^-4^) and probe periods (cluster-corrected *p* <= 2×10^-4^). (c) The topography of F-statistics averaged from 0 to 2.1 s after the onset of sample array indicated that the effect of set-size manipulation was clustered in parieto-occipital electrodes.

Next, we mapped parieto-occipital EEG activity into Holo-Hilbert spectral representations, then verified the load effect using the same ANOVA procedure as the above. The results were similar to previous studies (e.g., Proskovec et al., 2019). A sustained negative cluster was observed throughout the delay and probe periods in the alpha- and beta-bands (8-23 Hz; cluster-corrected *p* = 4×10^-4^; Fig. 4a). Compared with alpha power suppression, the effect of beta power (12-23 Hz) suppression was more pronounced in the probe period (1.1 to 2.1 s from the onset of sample array). On the other hand, transient positive clusters were observed in the theta-band (2-8 Hz) frequency during the onsets of the sample (cluster-corrected *p* = 6.2×10^-3^) and probe arrays (cluster-corrected *p* = 4×10^-4^). To further quantify the time course of inter-frequency power modulation in terms of AMF for the sustained negative cluster observed, we calculated the *AMF-t* spectrum within the alpha and beta time scales (i.e., IMFs 4 and 5), then performed the same analysis as for the *CF-t* spectrum. The results showed that the sustained negative cluster observed was consisted of two AMF components (cluster-corrected *p* = 5×10^-4^, Fig. 4b): a low-frequency component (0-2 Hz) extended throughout the task procedure, and a 2-12 Hz component that was only observed in the latter parts of the delay and probe periods. Note that the 2-12 Hz component was observed in the same time window in which the power of alpha CF gradually increased from the dips following the onsets of the sample and probe arrays (the blue trace in Fig. 4b). We also tested the load effect for frontal EEG power, but no significant effect was observed (Fig. S2). For comparison between HHSA and traditional time-frequency analysis, a separate analysis with decomposition using the Morlet wavelet transforms was illustrated in the Fig. S3. The patterns of the results were similar across the two methods. However, HHSA showed more precise time-frequency resolution, and only the HHSA could provide information in terms of AMF.

**Figure 4.**
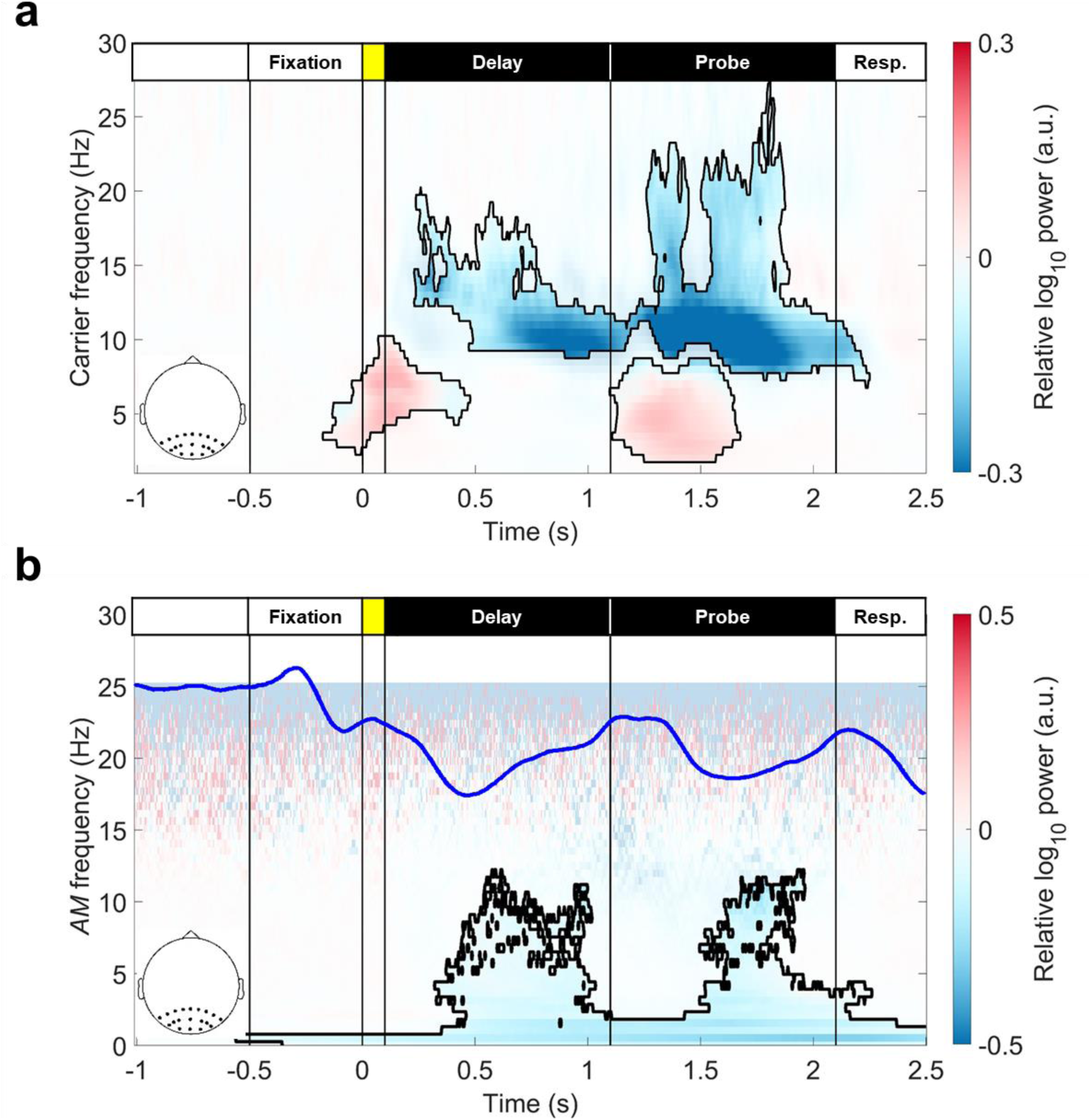
*CF-t* and *AMF-t* representations of load-dependent modulation of parieto-occipital EEG power. (a) Mean *CF-t* spectrum for set sizes 2, 4, and 6, contrasted with the set-size 1 condition. Load effects were verified by performing a repeated-measure ANOVA for each spectral point and were corrected by a cluster-based permutation procedure of 5000 iterations (both cluster-forming threshold and cluster significant level were set as *p* = 0.05). Black contoured areas denote significance after correction for multiple comparisons. Here, a sustained negative cluster was identified during the delay and probe periods (cluster-corrected *p* = 4×10^-4^), and two positive clusters could be observed after the onsets of the sample (cluster-corrected *p* = 6.2×10^-3^) and probe arrays (cluster-corrected *p* = 4×10^-4^). (b) The same analysis was performed for the *AMF-t* spectrum of IMFs 4 and 5 (6.4-23.6 Hz). Here, the results showed different time courses for different AMs of sustained load-dependent power suppression observed. The low-frequency AMF component (0-2 Hz) extended throughout the task procedure, whereas the 2-12 Hz AMF component only showed significance after processing the sample and probe stimuli (i.e., 0.5 s after stimuli onsets, cluster-corrected *p* = 5×10^-4^). The blue trace showed the fluctuation

### WM precision and HHS in the parieto-occipital area

After evaluating the load effects across different set sizes, we next conducted a correlation analysis for the individual differences between WM precision (κ) and parieto-occipital HHS. The set-size 1 served as the baseline condition. Both the κ value and HHS were subtracted from the one-item condition, and were transformed into z scores for analysis. The correlation was evaluated by a general linear model between the κ value and each spectral point of the HHS, with WM load as a covariate. The results were corrected with a cluster-based permutation test with 5000 iterations. The results for the *CF-t* spectrum showed a sustained negative correlate in alpha-band (9-12 Hz) for both the delay (cluster-corrected *p* = 0.007) and probe periods (cluster-corrected *p* = 0.002; Fig. 5a). Beta power (17-26 Hz) was negatively correlated with the κ value in the last 0.4 s time window of the probe period (cluster-corrected *p* = 0.013). In addition, theta power (3-7 Hz) also showed a negative correlation with the κ value toward the end of the probe period (the cluster was connected to the alpha-band cluster in the response period; cluster-corrected *p* = 0.002). The same analysis with the wavelet decomposed yield similar results (Fig. S4).

**Figure 5.**
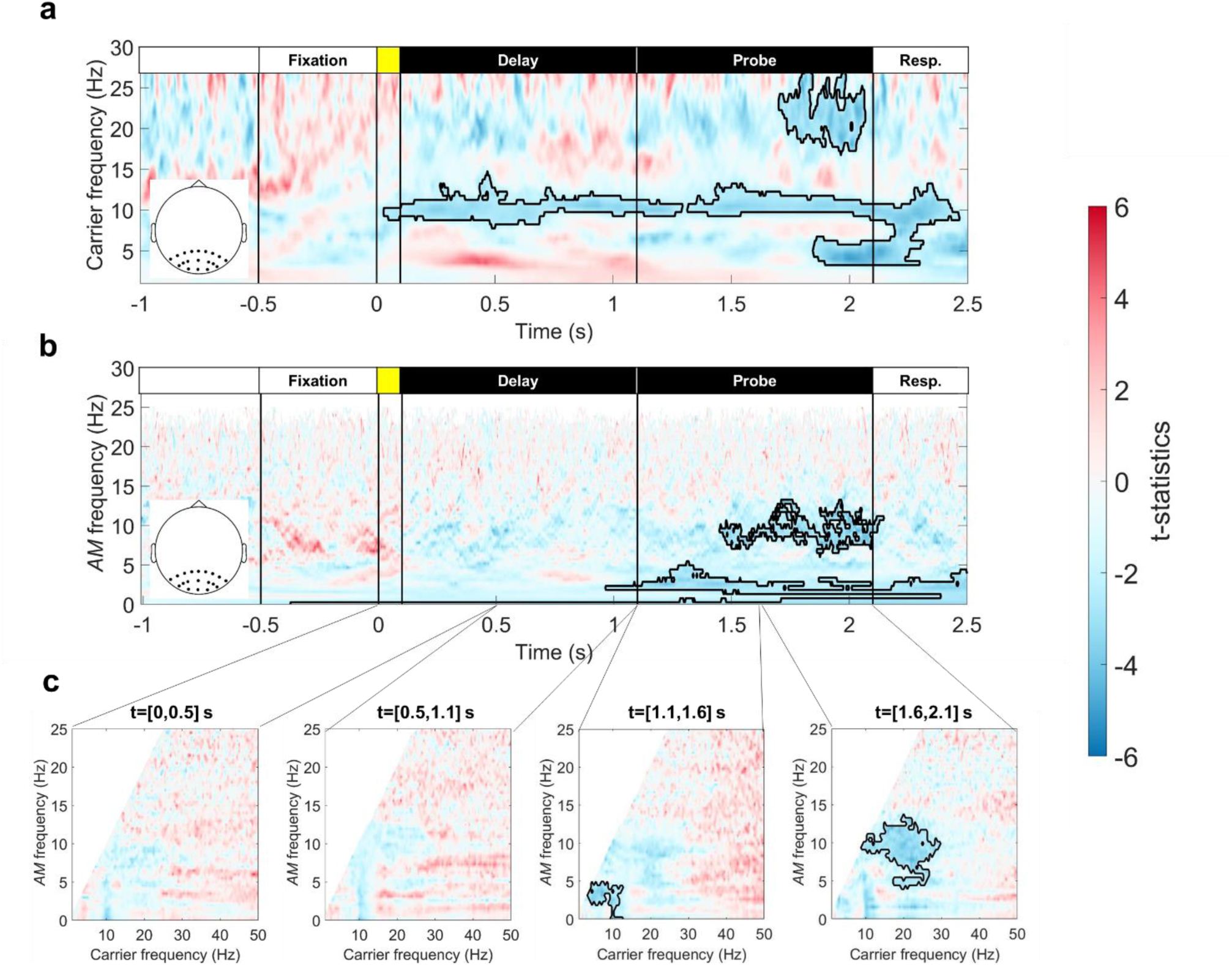
Representations of correlation strengths between parieto-occipital HHS and WM precision. Parieto-occipital alpha/beta power was negatively correlated to WM precision (κ). Black outlines denote significant correlation after cluster-based permutation correction (5000 iterations, *p* < 0.05 in a two-tailed test). (a) For the *CF-t* spectrum, a sustained negative correlation could be identified in the alpha-band (9-12 Hz) in the delay (cluster-corrected *p* = 0.007) and the probe periods cluster-corrected *p* = 0.002). In addition, a negative beta (17-26 Hz, cluster-corrected *p* = 0.013) and theta clusters (3-7 Hz, cluster-corrected *p* = 0.002) to κ value were also observed near the end of the probe period. (b) The *AMF-t* spectrum for the first-layer IMFs 4 and 5 (6.4-23.6 Hz) showed three AMF correlates to WM precision: an alpha-AMF correlate (7-13 Hz) was observed in the latter part of the probe period (cluster-corrected *p* = 0.0152), a delta-AMF correlate (1.5-4 Hz) was observed from the onset of probe array and was extended to the response period, and a low-frequency correlate (0-0.5 Hz) was observed throughout the task procedure (both belong to the same cluster, cluster-corrected *p* = 0.0044). (c) The results for *CF-AMF* spectra showed a negative correlation with WM precision during the probe period. In the first 500-ms time window of the probe period (t= [1.1, 1.6]), a significant cluster for theta/alpha CF was modulated in 2-5 Hz AMF (cluster-corrected *p* = 0.0106); in the last 500-ms time window (t= [1.6, 2.1]), a correlate of beta CF modulated in 5-12 Hz AMF (cluster-corrected *p* = 0.005) was identified.

For the *AMF-t* spectrum, three AMF correlates of WM precision could be identified (Fig. 5b). First, an alpha-AMF correlate (7-13 Hz) was observed in the latter part of the probe period (1.5 to 2.1 s from the onset of sample array; cluster-corrected *p* = 0. 0152). Second, a low-frequency AMF component (1.5-4 Hz) was observed from the onset of the probe array, and it extended to the response period. Finally, the trend component (0-0.5 Hz) showed sustained negative correlation throughout the task procedure. The latter two correlates were connected in the response period (cluster-corrected *p* = 0.0044). Further GLM analyses for the *CF-AMF* spectra showed that the parieto-occipital alpha/beta AM predicted individual WM precision after a target-defining cue was presented to the participants (Fig. 5c).

### WM precision and HHS in the frontal area

The GLM results for frontal HHS showed a positive correlation with individual WM precision (Fig. 6). For the *CF-t* spectrum, the theta-band power was positively correlated to κ value during WM retention. However, the frequency range of frontal correlates was slightly different in the delay and probe periods (Fig. 6a). The frequency range was 3-5 Hz for the cluster observed in the delay period (cluster-corrected *p* = 0.0054), whereas the frequency range was 5-8 Hz for the cluster observed in the probe period (cluster-corrected *p* = 0.0092). The comparable analysis with the Morlet wavelet transforms only showed a positive correlation trends but the statistics did not pass the significant level after multiple corrections (Fig. S5).

**Figure 6.**
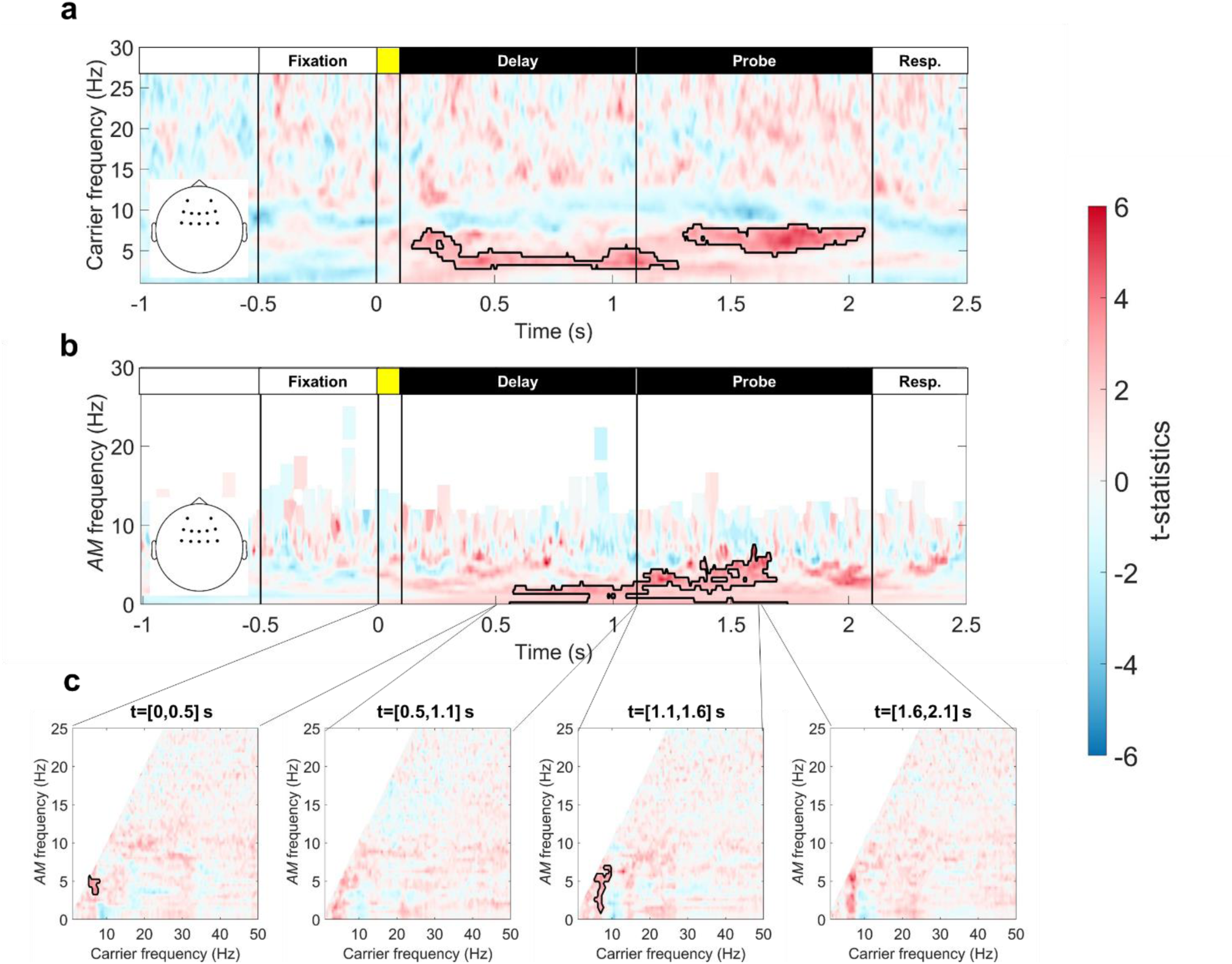
Representations of correlation strengths between frontal HHS and WM precision. Frontal theta power was positively correlated with WM precision (κ). Black outlines denote significant correlation after cluster-based permutation correction (5000 iterations, *p* < 0.05 in a two-tailed test). (A) The *CF-t* spectrum showed a positive correlation during WM retention, but the frequency of the correlate identified in the delay period was slower (3-5 Hz; cluster-corrected *p* = 0.0054) than the correlate identified in the probe period (5-8 Hz; cluster-corrected *p* = 0.0092). (B) The *AMF-t* spectrum for the first-layer IMFs 6 and 7 (1.5-11.8 Hz) showed a positive correlation with WM precision around the onset of the probe array (t = [0.5, 1.6] s). A significant positive cluster in the 1-4 Hz AMF range was observed during this time window (cluster-corrected *p* = 0.0074). (C) The *CF-AMF* spectra showed positive correlations with WM precision in the first 0.5-s time window after the sample array onset (modulated in 3-5 Hz AMF; cluster-corrected p = 0.0168) and the first 0.5-s time window after the probe array onset (modulated in 1-6 Hz AMF; cluster-corrected *p* =0.0076).

When the frontal theta power (IMFs 6 and 7, 1.5-11.8 Hz) was mapped into the *AMF-t* spectrum, a 1.5-4 Hz AMF component was observed in 0.5 s before and after the onset of the probe array (cluster-corrected *p* = 0.0074; Fig. 6). The result reflected the transition from the lower-theta (3-5 Hz) to the upper-theta (5-8 Hz) band as observed in Fig. 6a. The results of GLM analyses for the *CF-AMF* spectra in the same area showed that the AM component of the frontal theta power showed a significant correlation during the processing of incoming stimuli. A significant cluster could be observed in the first 500-ms time window of the delay (Fig. 6c, the first diagram; cluster-corrected p = 0.0168) and probe periods (Fig. 6c, the third diagram; cluster-corrected *p* = 0.0076).

### Intra- and inter-regional PACs between theta and alpha/beta oscillations

We revealed that the enhancement of frontal theta AM and the suppression of parieto-occipital alpha/beta AM predicted individual WM precisions. The next step is to test whether the local and inter-regional PACs change systematically according to the load manipulations or to the individual difference in WM precision. Under HHSA, the PAC between a slow and a fast oscillation can be calculated using the cross-frequency phase-locking values (cross-frequency PLV, Liang et al., 2021)Here, the phase of the slower oscillation was extracted from the first-layer IMFs, and the phase of the scale-matched AM of the faster oscillation was extracted from the second-layer IMFs. For this analysis, the PLVs between theta-, alpha-, and beta-band oscillations (i.e., IMFs 6, 5, and 4, respectively) were calculated across all electrode pairs, then were averaged according to set-size conditions, frequency combination, and the locations of electrode pairs. For comparison across WM loads, we found significant effects in both the first 0.5-s time windows after the onset of the sample array and that of the probe array (Fig. 7). During the onset of the sample array, the phase of frontal and parieto-occipital theta oscillations co-modulated parietal-occipital alpha power as set size increased (*F_(2,64_*_)_ = 7.3437, *p* = 1.35×10^-3^ for inter-areal coupling; *F_(2,64)_* = 11.453, *p* = 5.60×10^-5^ for intra-areal coupling). During the onset of the probe array, frontal and parieto-occipital theta oscillations not only modulated parieto-occipital alpha power (*F_(2,64)_* = 14.899, *p* = 4.87×10^-6^; *F_(2,64)_* = 22.406, *p* = 4.21×10^-8^), but also parieto-occipital beta power (*F_(2,64)_* = 7.187, *p* = 1.53×10^-3^; *F_(2,64)_* = 11.232, *p* = 6.59×10^-5^). The phase of parieto-occipital alpha oscillation also modulated beta power in a load-dependent manner (*F_(2,64)_* = 8.3092, *p* = 6.19×10^-4^). Finally, connectivity between frontal and parieto-occipital theta waves was also modulated by WM load during this period (*F_(2,64)_* = 5.4686, *p* = 6.42×10^-3^). All significant effects reported were positive, i.e., the PACs intensified with the increase of WM load. Details of the analyses were illustrated in the supplementary figure (Fig. S6).

**Figure 7.**
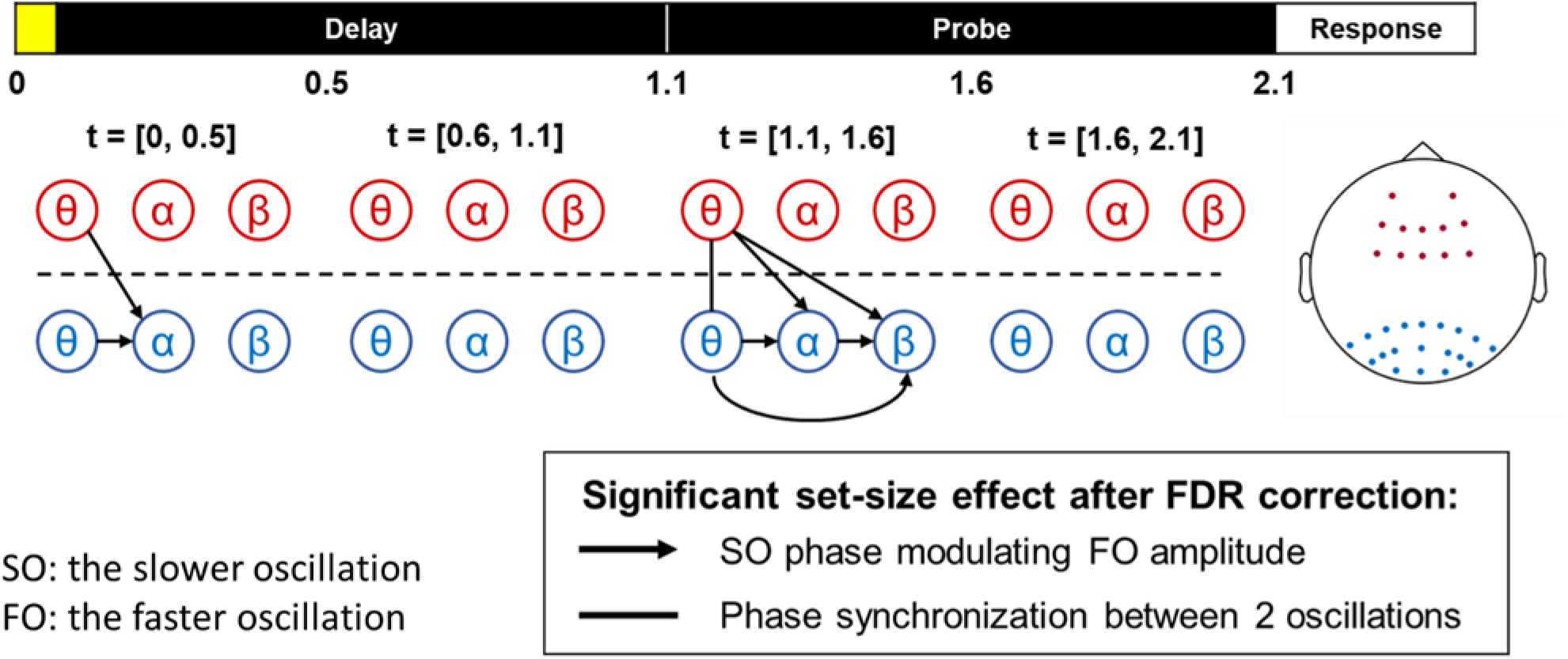
Load effects of intra- and inter-areal PLVs between frontal and parieto-occipital electrodes of interest. Nodes represent oscillatory activity in different timescales in frontal (red) and parieto-occipital (blue) electrodes of interest. Arrows between nodes indicate significant load effects (FDR < 0.05) in cross-frequency PLV. The arrow nock denotes the slower oscillation (SO) phase, and the arrowhead denotes the phase of the scale-matched AM of the faster oscillation (FO). A vertical line denotes a significant load effect in PLV between frontal and parieto-occipital oscillatory activities of the same timescale. Significant load effects were observed during the first 0.5-s following the onsets of the sample and probe array. All observed effects were positive. In the first interval (t = [0, 0.5] s), both local (within the parieto-occipital area) and long-range (between the frontal and parieto-occipital areas) PLVs for theta oscillation and the scale-matched alpha AM increased with set size. A similar pattern of results re-emerged and extended to the beta AM during the onset of the probe array (t = [1.1, 1.6] s). Furthermore, PLVs for frontal and parieto-occipital theta phases also increased during the onset of probe array.

We had performed similar GLM analyses between the individual differences in WM precision and PAC, but neither local nor interregional PACs predicted individual WM precision. Finally, the degree of PAC did not predict the subsequent suppression of the parieto-occipital alpha/beta AM. These results indicated that PAC and AM are independent measurements in WM processing.

## Discussion

This study investigated the electrophysiological bases of AM and PAC in WM processing. The main result showed that stronger load-dependent alpha/beta power suppression in parieto-occipital electrodes was associated with better WM precision. Furthermore, the observed sustained alpha/beta power suppression could be separated into continuous low-frequency (< 2 Hz) and 2-12 Hz AMF components. The low-frequency component extended throughout the task procedure, whereas the 2-12 Hz AMF component was observed only after processing the sample and probe stimuli. When correlating individual WM precision to alpha/beta AMs, we found negative correlations for the tonic alpha/beta power suppression (AMF <1 Hz) throughout the task procedure. On the other hand, higher AMFs of alpha/beta power suppression only showed significant correlations after the onset of the probe array.

Connectivity analysis with cross-frequency PLV showed a load-dependent increment of theta-alpha/beta PAC between the frontal and parieto-occipital areas and within the parieto-occipital area itself during the onsets of the sample and probe arrays. Therefore, the theta-AMF component of parieto-occipital alpha/beta AM was likely driven by frontal and parieto-occipital theta activity. The overall pattern of results suggests that WM precision, like capacity measurements, is supported by frontoparietal oscillatory activities (Liang et al., 2021; Palva et al., 2010; Siebenhühner et al., 2016).

### Alpha/beta power suppression reflects the ability to access feature information in WM

Our findings in parieto-occipital alpha/beta activity are consistent with previous reports showing that load-dependent alpha/beta suppression predicts individual differences in WM capacity (Erickson et al., 2019; Fukuda et al., 2015; Proskovec et al., 2019; Zammit et al., 2018) In the context of memory formation and retrieval, reducing alpha/beta power facilitates information-processing ability in task-relevant sensory areas (Hanslmayr et al., 2012). Therefore, the precision of memorized information is associated with the ability to maintain alpha/beta power suppression in areas storing stimulus-specific information (Griffiths et al., 2019). Since the κ value measured the quality of WM representation (Zhang and Luck, 2008), our results suggest that the maintenance of fine-grained perceptual information in WM requires engaging the parieto-occipital area (van Ede, 2018). This argument is consistent with studies using multivariate regression methods, which have shown that non-spatial feature information of visual WM could be decoded from bilateral posterior alpha/beta activities (Bae and Luck, 2018; Fukuda et al., 2016; de Vries et al., 2019). Similar results have also been reported in fMRI studies, in which the stimulus information can be decoded from the pattern of BOLD responses in visual areas (Ester et al., 2009; Harrison &Tong, 2009; Pratte and Tong, 2014). In addition, the same pattern of results can extend to other modalities, e.g., the load-dependent alpha/beta power suppression could either be observed within the visual or somatosensory area, depending on which modality was prompted for a response in an intermodal WM task (van Ede et al., 2017).

By decomposing parieto-occipital alpha/beta AMs into different AMFs, we found differences between the low-frequency component (<1 Hz) and higher AMF components (2-12 Hz) in their correlation to κ values. Specifically, the low-frequency component showed sustained negative correlation throughout the maintenance period, whereas higher AMF components only showed a significant correlation after the onset of the probe array. Furthermore, different CF-AMF combinations were observed during the processing of probe stimuli (the first 0.5-s time window after the onset of the probe array; t = [1.1, 1.6] s) and its later maintenance (t = [1.6-2.1] s; Fig. 4C). Alpha AM in the delta/theta band (2-5 Hz) predicted individual WM precision during the target selection period, whereas beta AM in the theta/alpha band (5-12) Hz showed a significant correlation in the later time window. These results might imply different cognitive factors contributing to the maintenance of WM information. The low-frequency component reflected tonic psychophysiological states such as alertness and sustained attention (Oken et al., 2006). In contrast, alpha AM in the delta/theta band reflected the target selection process (de Vries et al., 2018). One recent study has shown that the load-dependent increment of the speed of parietal alpha/beta oscillation predicts individual WM capacity (Noguchi and Kakigi, 2020). Therefore, the beta AM observed might indicate a more efficient target selection for high-precision participants.

### Relationship between frontal theta power and WM precision

Frontal midline theta activity has been considered the oscillatory substrate for cognitive control or cognitive efforts in completing the task (Cavanagh and Frank, 2014; McFerren et al., 2021). Prior studies have associated the increment of frontal midline theta power to increasing memory load (Jensen and Tesche, 2002; van Ede et al., 2017; Popov et al., 2018), increasing task difficulty (Wisniewski et al., 2017), or as an index of successful WM manipulation (Itthipuripat et al., 2013). The increment of frontal theta power is also associated with better performance in WM tasks (Gärtner et al., 2014; Zakrzewska and Brzezicka, 2014). Our results showed a positive correlation between frontal theta power and WM precision, however, frontal theta power did not scale with WM loads (Figs. S2 and 6A). The seeming contradiction of load effects could be explained as the following. First, frontal theta power increment is more sensitive in maintaining temporal order information than visual item information (Hsieh et al., 2011; Liu et al., 2020). Therefore, the effect size of frontal theta power increment would be smaller for concurrent visuospatial WM tasks. Second, other factors such as individual WM capacity and acute stress could interact with the frontal theta load effect, so that frontal theta power peaked at some critical load and then decreased at higher loads (Gärtner et al., 2014; Zakrzewska and Brzezicka, 2014). One recent intracranial EEG study also failed to replicate the load-dependent theta power increment in dACC, which was considered as the main source of frontal midline theta activity (Brzezicka et al., 2019; Meltzer et al., 2007). The study argued that a portion of the frontal midline activity observed from the scalp was indeed contributed by the hippocampus. Studies with frontal or frontoparietal non-invasive brain stimulations have shown modulations of WM performance when stimulating in the theta-band frequency (Alekseichuk et al., 2016; Reinhart and Nguyen, 2019; Riddle et al., 2020; Sahu and Tseng, 2021). Furthermore, the improvement of WM performance is associated with enhanced theta connectivity between frontal and posterior areas, indicating more efficient control for the information stored in WM (Reinhart and Nguyen, 2019; Sahu and Tseng, 2021)

### Frontoparietal cross-frequency PLV indicates the top-down modulation of memorized contents

Literature has suggested that the prefrontal cortex serves as the central executive, which monitors the overall task sequence rather than maintenance in WM (D’Esposito &Postle, 2015; Sreenivasan et al., 2014). In particular, the prefrontal cortex exerts control over posterior areas storing WM contents through low-frequency oscillations around the theta-band ( Fell and Axmacher, 2011; Herweg et al., 2020; Sauseng et al., 2010; Staresina and Wimber, 2019; de Vries et al., 2020). Our results support the idea by showing the load-dependent increment of cross-frequency PLVs (i.e., theta-alpha PAC) between the frontal theta phase and parieto-occipital alpha/beta power (Fig. 7). Furthermore, such long-range modulation is only transiently involved during the processing of incoming stimuli. The long-range interaction through PAC is consistent with a variety of recent works. First, previous works have shown that frontal-to-parietal theta synchronization and parietal-to-frontal alpha synchronization are crucial for successful memory retrieval (Johnson et al., 2017; Popov et al., 2018). Second, the local and long-range PAC are enhanced by memory demand, either between theta-alpha (de Vries et al., 2018) or theta-beta bands (Daume et al., 2017). In another study with a similar HHSA approach, the improvement of WM capacity from transcranial direct current stimulation is closely related to the inter-areal PAC between the frontoparietal theta phase and beta/gamma powers (Liang et al., 2021). Third, the transient modulation of posterior alpha activity by frontal theta oscillation has been reported in WM studies involving spatial cueing and multiple regression analysis (deVries et al., 2019; Wallis et al., 2015). This line of works often reported stronger alpha power suppression contralateral to the cued hemifield after cue-induced frontal theta power enhancement. Taken altogether, the pattern of long-range PAC observed reflects the online control of WM representations according to the task relevance at the moment.

### HHSA reveals the relationship between oscillation power variations and PAC

While conventional PAC methods calculate the distribution of the power of a high-frequency oscillation to the phase of a low-frequency oscillation, the current study isolates the estimation of cross-frequency coupling in two steps: first, the amplitude of an oscillation was decomposed into different AMs with HHSA; second, PAC was evaluated by cross-frequency PLV between the slow oscillation and the scale-matched AM of the fast oscillation (Juan et al., 2021; Liang et al., 2021). We have shown that WM load manipulation modulates theta-alpha and theta-beta PACs during the processing of probe stimuli, then was followed by a corresponding change in alpha/beta power suppression. However, only alpha/beta AM predicts individual WM precision. The results imply PAC and AM have distinct meanings in WM processing. Theta-alpha/beta PAC reflects the mechanism of top-down selection over WM representations (de Vries et al., 2020), whereas alpha/beta AM reflects the fidelity of target information after selection.

## Conclusion

Our results demonstrate that rhythmic modulations of parieto-occipital alpha/beta power during the retention period predict the recall precision of visual WM. AMF components of alpha/beta power reflect cognitive factors acting at different frequencies to modulate the precision of WM representations. The power of AMF components does not depend on any specific lower frequency. Results of cross-frequency PLVs between theta phase and alpha/beta AM reflect the online interaction of oscillations across different frequencies for the temporal involvement of different cognitive processes in WM.

## Materials and Methods

### Participants

Thirty-three university students were recruited to participate in the experiment (21 males and 12 females, mean age = 23.13 years and standard deviation = 5.16 years). All participants had normal or corrected-to-normal vision and all gave informed consent before participation. The study procedures were approved by the local IRB committee (National Taiwan University Research Ethics Committee, 201606ES031).

### Experimental design

The behavioral task was modified from the color recall tasks for WM precision (Bays et al., 2009; Zhang and Luck, 2008). The stimuli were presented on an LCD monitor in a black background at a viewing distance of 60 cm. The colors were selected from a circle in CIELAB space (Robertson, 1977), centered at L=70, a=20, b=38, in CIE coordinates with a radius of 60. The items to be remembered were a set of color squares with size 2°x2° of visual angle. The color of each item was sampled from 45 possible color values at every 8° of the circle. The locations of the items were randomly chosen but equally spaced concentrically, 4.5° away from the center of the monitor. The color wheel for participants to use to make their selection was constructed with 360 color values, sampled from 1° to 360° of the color circle. The color wheel was centered on the screen, with an inner and outer radius of 6° and 8.2° of visual angle.

Each trial began with a concurrent presentation of a beep sound (500 Hz) and a central fixation cross (gray, 1°×1°). The beep sound lasted for 100 ms, and the fixation cross remained on display throughout the trial. After 500 ms, a set of color squares was displayed for 100 ms, followed by a 1000-ms delay period. After the delay period, a set of outlined squares was presented at the location of each item during stimulus presentation for 1 s (the probe period). One of the squares had thicker edges, indicating the target item for the observer to recall. A color wheel was displayed as the probe squares vanished. The observer had to identify the target color by mouse-clicking the corresponding color in the color wheel. During the response period, the color in the target location kept updating to the color selected by the mouse cursor (Fig. 2a). The subsequent trial began 500 ms after a target color was picked from the color wheel. In different experimental conditions, the set size of the sample array varied between 1, 2, 4, or 6 items. The four set-size conditions were randomly interleaved across trials. All participants performed 90 trials for each condition, making 360 trials in total.

### Mixture model analysis

Behavioral performance was assessed using mixture model analysis (Bays et al., 2009). In each trial, the degree of error was obtained by calculating the angular deviation between the coordinates of the reported color and the correct target color in the color wheel. The model proposed three sources of WM errors: 1) the probability that the target color was not stored in the WM system, 2) the probability of misreporting a non-target color instead of the target color, and 3) information was stored with noise in WM. Participants were assumed to make a uniformly random response when they could not recall the target color. In addition, all items were assumed to be encoded with equal resolution and encoding to decay with equal rate. The probability of the reported color value in each trial can therefore be described as follows:

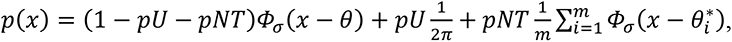

where *Φ_σ_* represented the von Mises (circular Gaussian) distribution with mean zero and standard deviation *σ*, *x* was the reported color value, *θ* was the target color value, 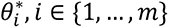 were the *m* non-target color values, *pT* was the probability of correctly remembering the target color, *pU* was the probability of random guessing, and *pNT* was the probability of misremembering the target location (Fig. 2b). The index for WM precision (*κ* = 1/*σ*^2^) was the concentration number of the von Mises distribution, which was the reciprocal of error variance.

### Recording and preprocessing of EEG data

EEG was continuously recorded throughout the experiment using SynAmps2 amplifiers (Compumedics Neuroscan) using Ag/AgCl electrodes mounted in a 64-channel electrode cap (Quik-Cap, Neuromedical Supplies). The EEG cap followed the international 10-20 system for electrode placement. Impedances of all electrodes were kept below 5 kΩ. The sampling rate was set at 2000 Hz, and horizontal and vertical electrooculograms were also recorded along with the EEG recordings. The recordings were filtered with a low-pass filter of 500 Hz, and AFz served as the ground electrode.

The overview of EEG data analysis was illustrated in Fig. 8. For preprocessing, the raw EEG data were offline-referenced to the average of M1 and M2 electrodes. The data were epoched from -1500 to 3000 ms relative to the onset of the sample array. Epochs containing artifacts of muscle activity, head or body movements, and electrode noise were rejected by visual inspection. Eye movements and blink artifacts were identified and removed with independent component analysis (ICA) using the EEGLAB toolbox (Delorme and Makeig, 2004). Epochs deviating more than 200 μV were also rejected. Electrodes containing more than 50% of such ‘bad’ trials were replaced by spline interpolation of their neighbors. 19 out of 33 participants required such interpolation (three channels for one participant, two channels for four participants, and one channel for 14 participants). Most such electrodes were prefrontal or temporal electrodes (FP1/2, FPz, T7/8, TP7/8, and C6). After this procedure, none of the participants was rejected for further analysis. Finally, the preprocessed EEG signal was transformed into its scalp current sources via the surface Laplacian method (Perrin et al., 1989). The application of surface Laplacian not only highlighted the local features, but also reduced the volume-conduction effects across electrodes. The transformed EEG signal was decomposed with HHSA, and the PAC between different frequency components was estimated with cross-frequency PLV.

**Figure 8.**
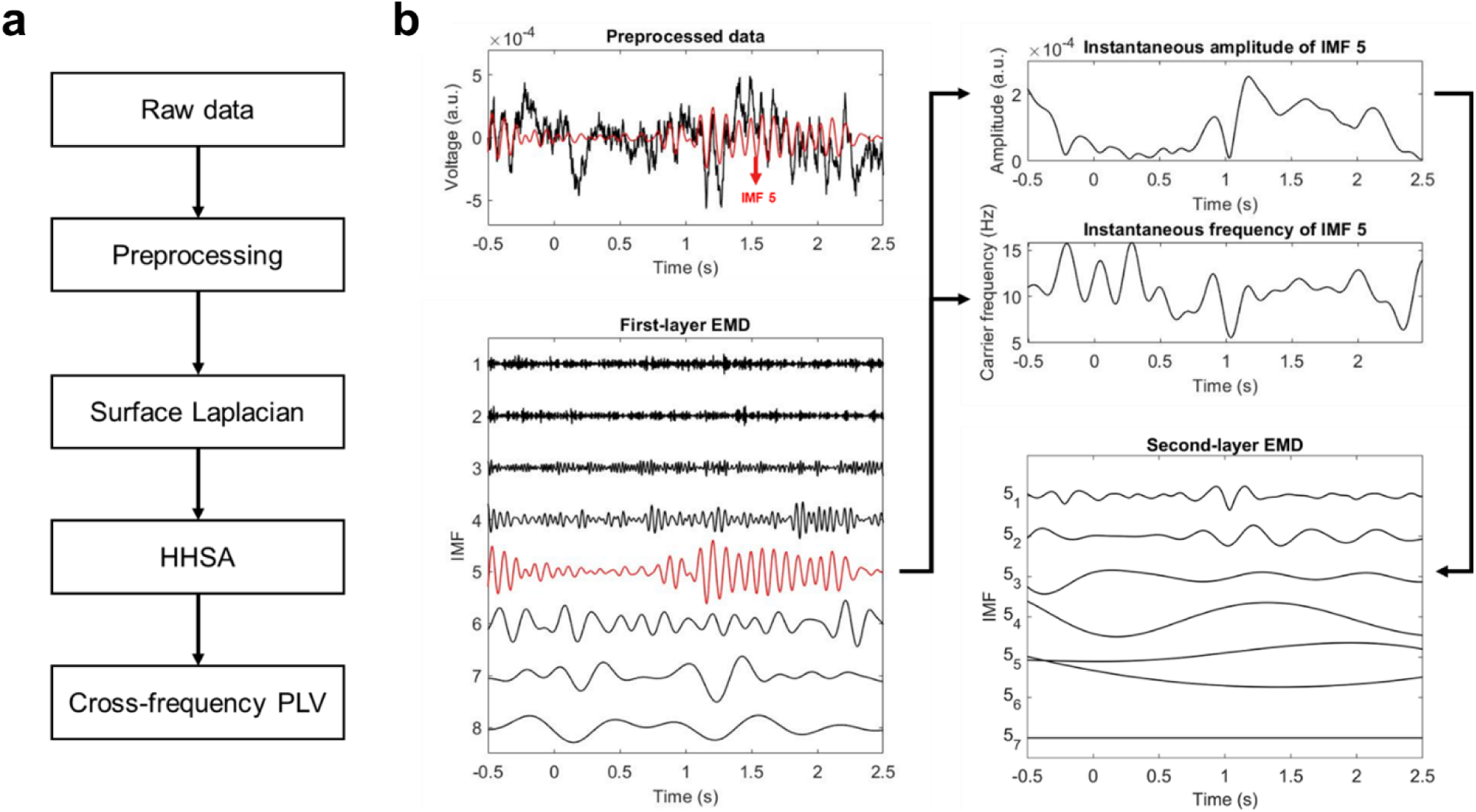
The overview of EEG data analysis. (a) The pipeline of EEG data analysis. EEG recordings was preprocessed with regular procedures as described in the main text. The preprocessed data was mapped into its scalp current density using the surface Laplacian, then was subjected to HHSA. The phase of a slow oscillation, and the phase of frequency-matching AM power of a faster oscillation could be directly obtained with HHSA. Therefore, the PAC amplitude between two oscillations could be estimated with statistics such as PLV. (b) The procedure of HHSA involved two layers of EMD. The input signal was decomposed with the first-layer EMD to extract the oscillations of interest, for instance, the alpha-band oscillation (marked in red). The instantaneous frequency and envelope (i.e., AM) of IMFs could be calculated in direct quadrature. The envelope of the interested oscillation was further decomposed with the second-layer EMD, yielding another set of IMFs. The second-layer IMFs represented different modes of AM oscillations of the envelope.

### Holo-Hilbert Spectral Analysis (HHSA)

EEG data were analyzed by HHSA (Huang et al., 2016; Juan et al., 2021; Liang et al., 2021; Nguyen et al., 2019) with in-house Matlab code. The method is based on two-layers of empirical mode decomposition (EMD) and a high-dimensional spectral representation. EMD is a process that decomposes data into a set of intrinsic mode functions (IMF, Huang et al., 1998) from fastest to slowest oscillatory timescales. Since each IMF is narrowband and symmetric locally with a zero mean, it is ideal for estimating instantaneous oscillatory phases with Hilbert transform or later proposed methods such as normalized Hilbert transform and direct quadrature (DQ, Huang et al., 2009)

Transient bursts in biological signals sometimes cause mode mixing, i.e., a single IMF contains multiple frequency components with classical EMD. Here, we applied an improved version of EMD utilizing sinusoidal masking signals in conjunction with the original EMD algorithm to reduce the problem (Deering and Kaiser, 2005; Nguyen et al., 2019; Quinn et al., 2021). A set of masking signals with initial phases 0, π/2, π, and 3π/2 were added into a pre-decomposed IMF obtained from regular EMD to enhance decomposition. The optimal amplitude of the masking signals is 1.6 times above the averaged amplitude of the components through experience (Deering and Kaiser, 2005). We used a factor of 2 for both layers of EMD. The frequency of masking signals was the same as the mean instantaneous frequency of the pre-decomposed IMF. Instantaneous frequencies of IMFs were directly calculated by taking time derivatives of instantaneous phases obtained from DQ.

In HHSA, the first-layer IMFs were obtained by performing EMD on the preprocessed data (Fig. 8B). The second-layer IMFs were obtained by performing the same EMD to the envelope functions of the first-layer IMFs. To distinguish the instantaneous frequencies of the first- and second-layer IMFs, we will refer to instantaneous frequencies of the first-layer IMFs as “carrier frequency” (CF) and to second-layer IMFs as “AM frequency” (AMF). AMF represented rhythmic variations of the slow-changing envelope resulting from all-important cross-scale interactions (Huang et al., 2016; Liang et al., 2021). Compared to conventional approaches, HHSA has the following advantages: 1) it is immune to the uncertainty principle of time-frequency analysis (i.e., the trade-off between time and frequency resolutions) by providing instantaneous frequency (Huang et al., 1998, 2009), so the data can be represented in high precision in the spectral domain; 2) the structure of data itself determines the timescales being examined, so *a priori* assumptions for frequency bands are not needed; 3) the envelope function of each frequency component can be decomposed again into a set of second-layer IMFs, where each second-layer IMF represents modulation of the envelope function within a dyadic timescale.

The result of the two-layer EMD was further represented by a 3D *Holo-Hilbert spectrum* (HHS) with time (*t*), CF, and AMF. In practice, HHS is projected along either of the three dimensions to highlight different aspects of the full spectrum. Projecting the HHS onto the *CF-time* plane created the *CF-t* spectrum (Fig. 9a), which captured the intra-mode frequency variation of the carriers. *CF-t* spectrum was equivalent to the spectrogram obtained from Hilbert-Huang Transform (HHT, Huang et al., 1998) that has been frequently used for analyzing the instantaneous frequency and amplitude variations of biological signals (e.g., Lopes-dos-Santos et al., 2018; Quinn et al., 2021). When HHS was projected onto the *AMF-time* plane, the result highlighted the dynamic of cross-scale interaction for a given range of carrier frequencies (the *AMF-t* spectrum, Fig. 9b). When HHS was projected onto the *AMF-CF* plane, the results showed the overall structure of cross-scale interactions between different frequency components in a given period (the *AMF-CF* spectrum, Fig. 9c). In this study, the resolution of HHS was set as 4 ms for time and 0.5 Hz for both CF and AMF.

**Figure 9.**
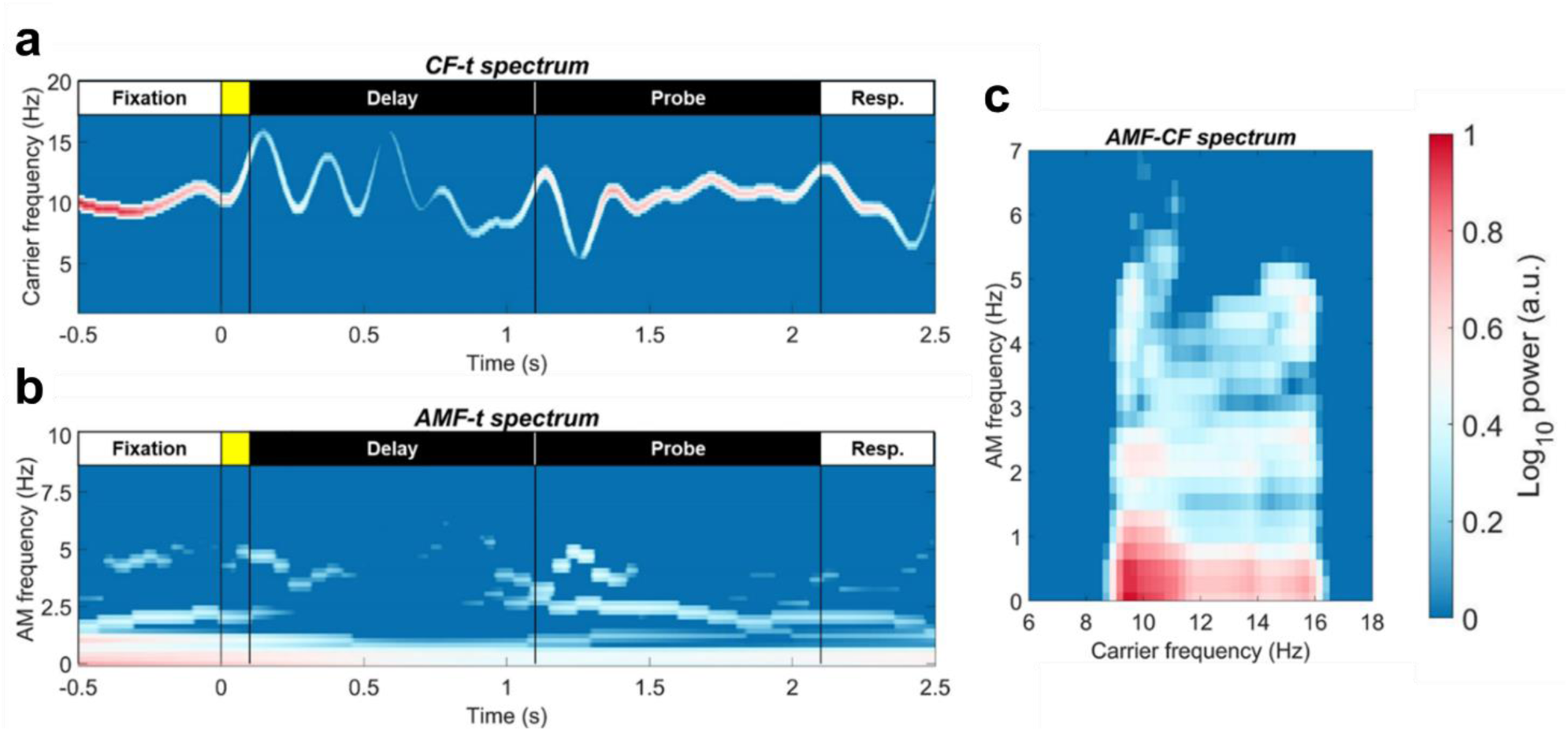
An illustrative example of spectral representations using HHSA. The example is generated with the alpha-band IMF as illustrated in Fig. 8. Each spectrum highlights different characteristic of the signal. (a) The *CF-t* spectrum shows the power and frequency variations of the alpha-band IMF over time. It is similar to spectrograms obtained using classical methods (e.g., wavelet transforms), but with higher time-frequency resolutions. (b) The *AMF-t* spectrum shows the AM power at various modulating frequencies over time. (c) The *AMF-CF* spectrum shows the power distribution of carrier frequencies and their AM (in different modulating frequencies) within a given period.

### Intra- and inter-areal PAC under the HHSA framework

IMF is a function that requires (1) symmetric locally to zero, and (2) there exists only one extremum between two successive zero-crossing points. According to the above definition, the instantaneous phases for the first- and second-layer IMFs could be obtained directly with DQ. With the phase functions of carriers and their AMs given, the degree of PAC could be measured by phase synchronization indices such as phase-locking value (PLV):

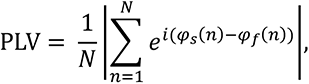

in which *φ_s_* and *φ_f_* denote the phase functions of two oscillations, and *N* denotes the number of sampling points within a time window. When PAC employs the PLV formula under the HHSA framework, one phase function is from the carrier of a slower oscillation, and the other is from the scale-matched AM component of a faster oscillation. This measure of PAC in the HHSA framework is referred to as Holo-Hilbert cross-frequency phase clustering (HHCFPC) and has been demonstrated as efficient in quantifying intra- and inter-areal CFC in the process of WM (Liang et al., 2021).

### Statistical analyses of EEG data

We used Matlab and the FieldTrip Analysis Toolbox for statistical analysis of EEG data (Oostenveld et al., 2011). Epochs were time-locked to the onset of the sample array for all analyses.

### Evaluating the effects of WM load on EEG activity

For preliminary analysis, we calculated the mean amplitude of each IMF across all electrodes for each set size and participant, then tested the effect of load manipulation with a repeated-measures ANOVA for set sizes 2, 4, and 6 at each time point. The single-item condition served as active control for the task. ANOVA results were corrected for multiple comparisons by a non-parametric cluster-based permutation test of 5000 iterations. The method only assumes that every experimental condition is associated with a probability distribution. Unlike parametric tests, the validity of the permutation test does not depend on the probability distribution of the data nor the cluster-forming method (Maris and Oostenveld, 2007). Both cluster-forming threshold and the critical value of significance were set at *p* = .05.

Based on previous studies (Erickson et al., 2019; Fukuda et al., 2015; de Vries et al., 2018), we choose mid-frontal electrodes (AF3/4, F1/2/3/4, FC1/2/3/4, Fz, and FCz) and parieto-occipital electrodes (P1/2/3/4/5/6/7/8, PO3/4/5/6/7/8, O1/2, Pz, POz, and Oz) for area-based analysis. Averaged *CF-t* power spectra in frontal and parieto-occipital electrodes were calculated separately, ranging from 1 to 50 Hz. Load effects for the *CF-t* spectra were evaluated with the same procedure of repeated-measures ANOVA as mentioned. If any significant cluster was identified, the *AMF-t* spectrum of the corresponding CF range was calculated and tested with the same ANOVA procedure.

### Correlation between individual WM precision and EEG activity

The same *CF-t* and *AMF-t* representations of parieto-occipital EEG power were also used for correlation analyses. EEG spectra and *κ* values from the set-size one condition were subtracted from those from set sizes 2, 4, and 6, and were then transformed into z scores for regression analysis. For each spectrum, a general-linear model (GLM) was constructed in which the z-transformed *κ* values served as a regressor to predict the z-transformed EEG power in each spectral point. Since both κ values and EEG power varied systematically with increasing set size, the regression weights were separately evaluated for set sizes 2, 4, and 6, then averaged together for statistical tests (i.e., set size served as a control variable for the GLM). The approach was similar to previous within-subject studies (e.g., Myers et al., 2014; Poliakov et al., 2014) but on a between-subject basis. The same cluster-based permutation procedure was applied to correct the results of GLM as for load effects, but both cluster-forming threshold and critical value were set at *p* = .05 for two-tailed tests. In addition, we also tested EEG activity in frontal electrodes to see if there was any WM correlate in the cluster. Correlations between frontal *CF-t* spectrum and WM precision were analyzed first; if any significant correlate was identified, the *AMF-t* spectrum of the corresponding CF range was subjected to analysis.

### Intra- and inter-areal PAC

EEG data were decomposed into first layer IMFs via EMD. In this dataset, IMFs 4, 5, and 6 represented oscillatory activities in beta-, alpha-, and theta-band, respectively (Fig. 2a). We calculated cross-frequency PLVs between IMFs 4, 5, and 6 across all possible electrode pairs, then averaged PLVs within and between the frontal and parieto-occipital electrodes of interest for each set-size condition. A repeated-measures ANOVA was applied for set sizes 2, 4, and 6 to test the effect of load manipulation with the one-item condition serving as the control. Time windows for PLV evaluation were determined according to the previous results of load effects over HHS (Fig. 3b). The first interval ranged from 0 to 0.5 s after the onset of the sample array, and the remaining part of the delay period constituted the second interval. Time windows for the probe period were also divided into the first 0.5 s and the remainder interval for analysis. The results were corrected for multiple testing using the Benjamini-Hochberg procedure (Benjamini and Hochberg, 1995) with a false-discovery rate (FDR) of 0.05.

## Acknowledgments

This work was funded by the Ministry of Science and Technology, Taiwan (grant numbers:110-2639-H-008-001-ASP; 109-2410-H-008-044-; 109-2321-B-037-001-; 108-2321-B-075-004-MY2; 110-2410-H-008-039-; 110-2410-H-008-040-MY3) and Taiwan Ministry of Education’s “Academic Strategic Alliance: Taiwan and Oxford University” project grant (MOE Oxford-NCU-CIPH collaborative project). We are grateful to Dr. Shih-kuen Cheng for sharing his expertise on EEG and the recording equipment.

---

**Figure S1.**
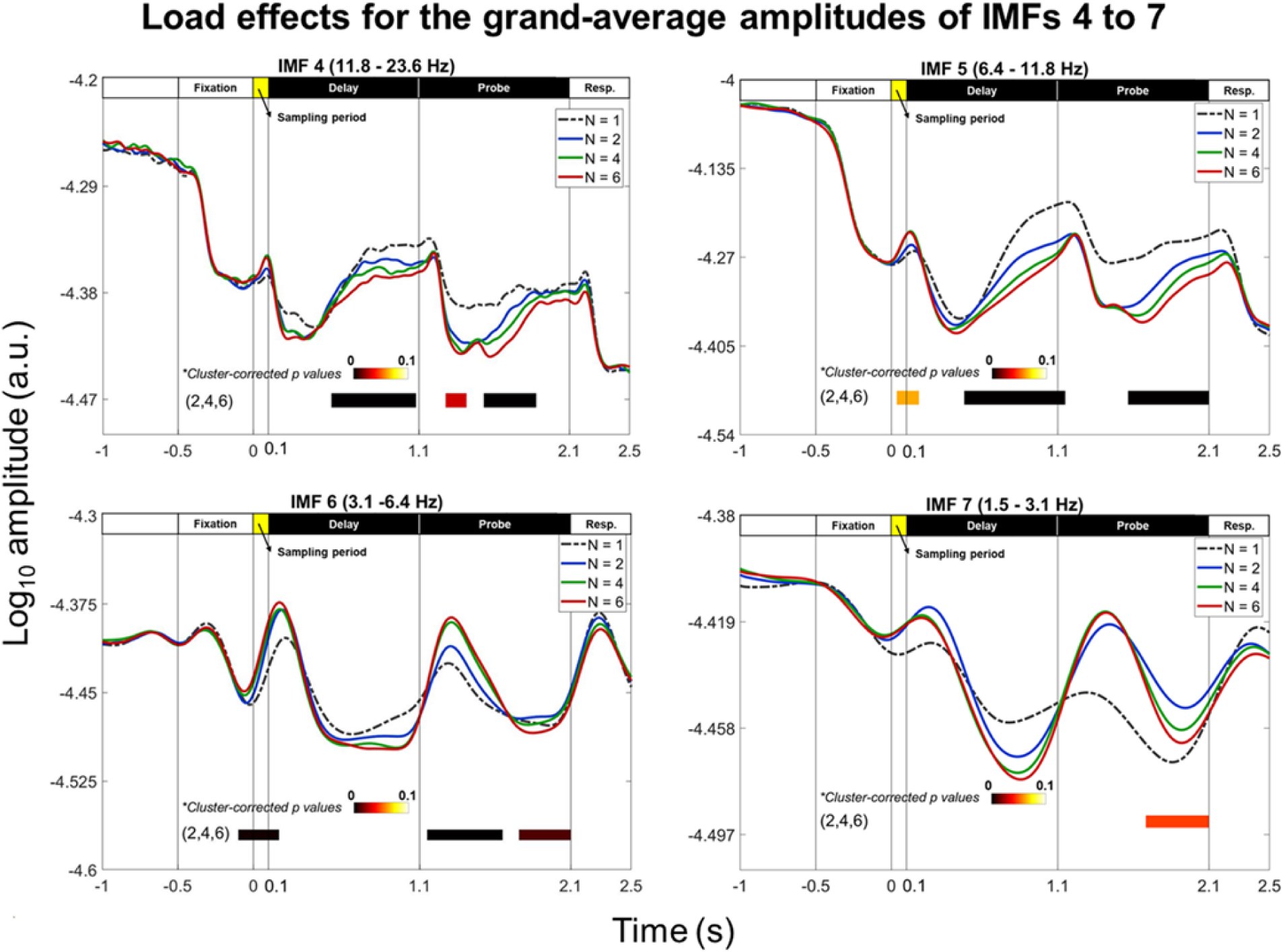
Load-dependent modulation of powers in IMFs 4—7. A repeated-measure ANOVA between set sizes 2, 4, and 6 was performed for each time point and each IMF with the single-item condition served as the baseline for comparison. Results of ANOVA were corrected by a cluster-based permutation correction for 5000 iterations. Color bar denoted cluster-corrected *p*-value for each cluster. (*upper panel*) IMFs 4 and 5 showed similar negative load effects after processing the sample and probe stimuli, as indicated by the transient power increment following the onset of the sample and probe arrays. IMF 5 also showed a transient positive load effect following the onset of the sample array. (*lower panel*) IMF 6 showed transient positive effects following the onset of the sample and probe arrays, and both IMFs 6 and 7 showed negative load effects in the last 0.4-s time window of the probe period.

**Figure S2.**
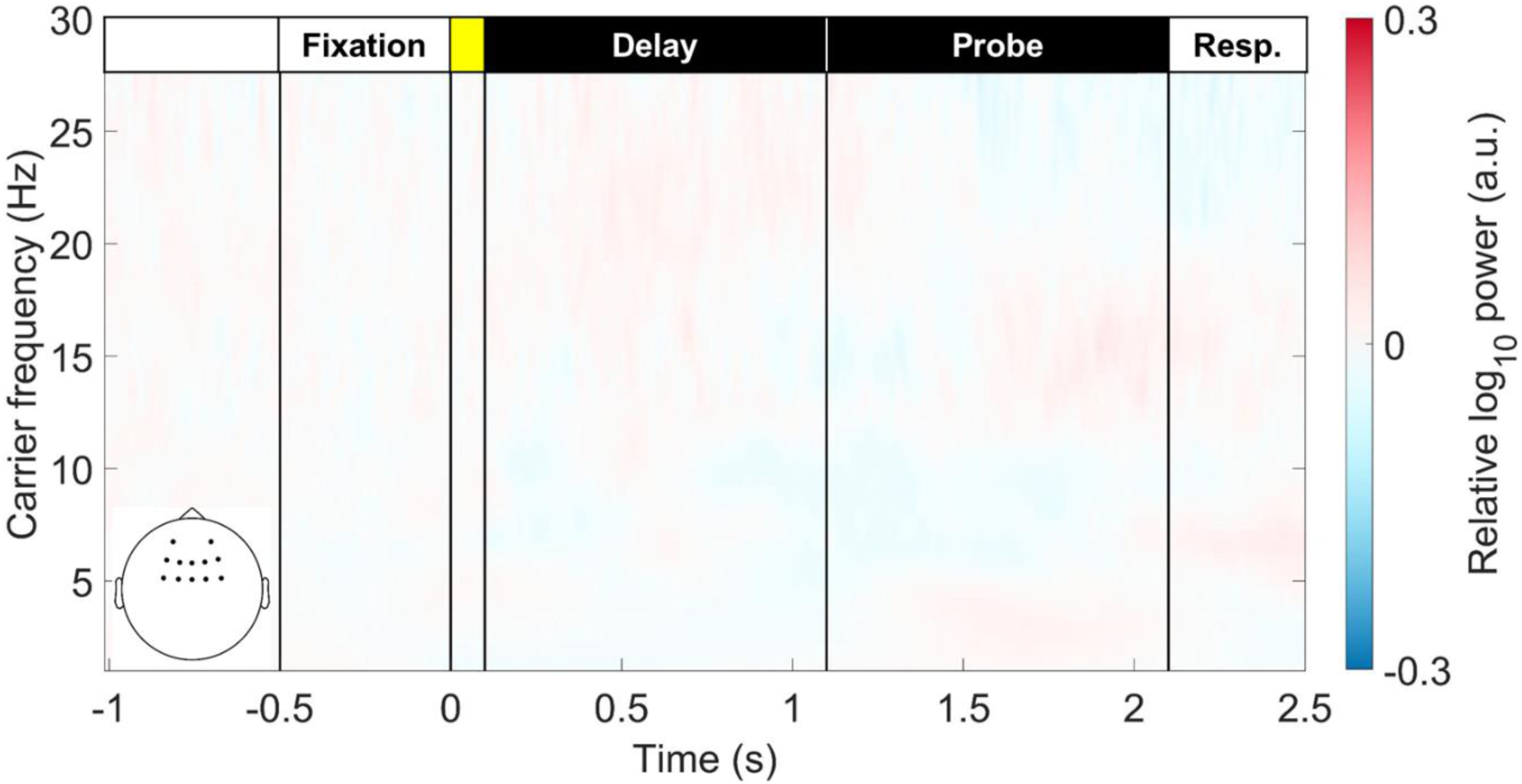
The *CF-t* representation of load effects for frontal EEG power. The figure illustrated the averaged *CF-t* spectrum in the frontal area for set sizes 2, 4, and 6 contrasting with the single item condition. No significant load effect was detected throughout the WM retention period after cluster-based permutation correction (both cluster-forming and significance thresholds were set at *p* = 0.05).

**Figure S3.**
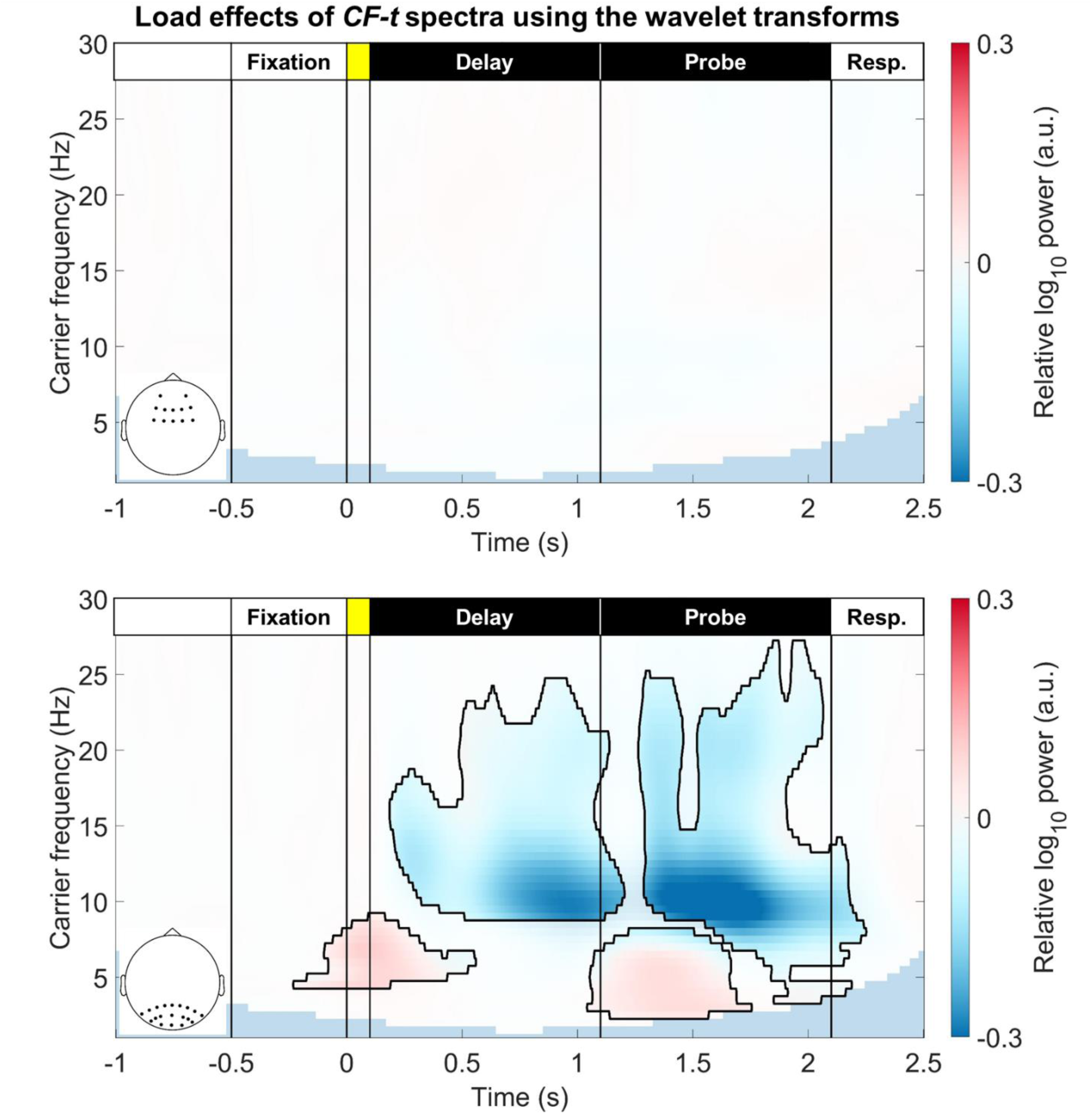
The *CF-t* representation of load effects for frontal and parieto-occipital EEG using the Morlet wavelet transforms. The effects of load manipulation were similar in HHS and the wavelet-based *CF-t* spectra. No effect was observed in the frontal region. Positive theta-band clusters could be observed in the onset of the sample (cluster-corrected *p* = 6.2×10^-3^) and probe (cluster-corrected *p* = 6.2×10^-3^) arrays. Negative clusters in the alpha and beta bands were observed in the delay (cluster-corrected *p* = 6.2×10^-3^) and probe (cluster-corrected *p* = 6.2×10^-3^) periods, after the onsets of stimuli.

**Figure S4.**
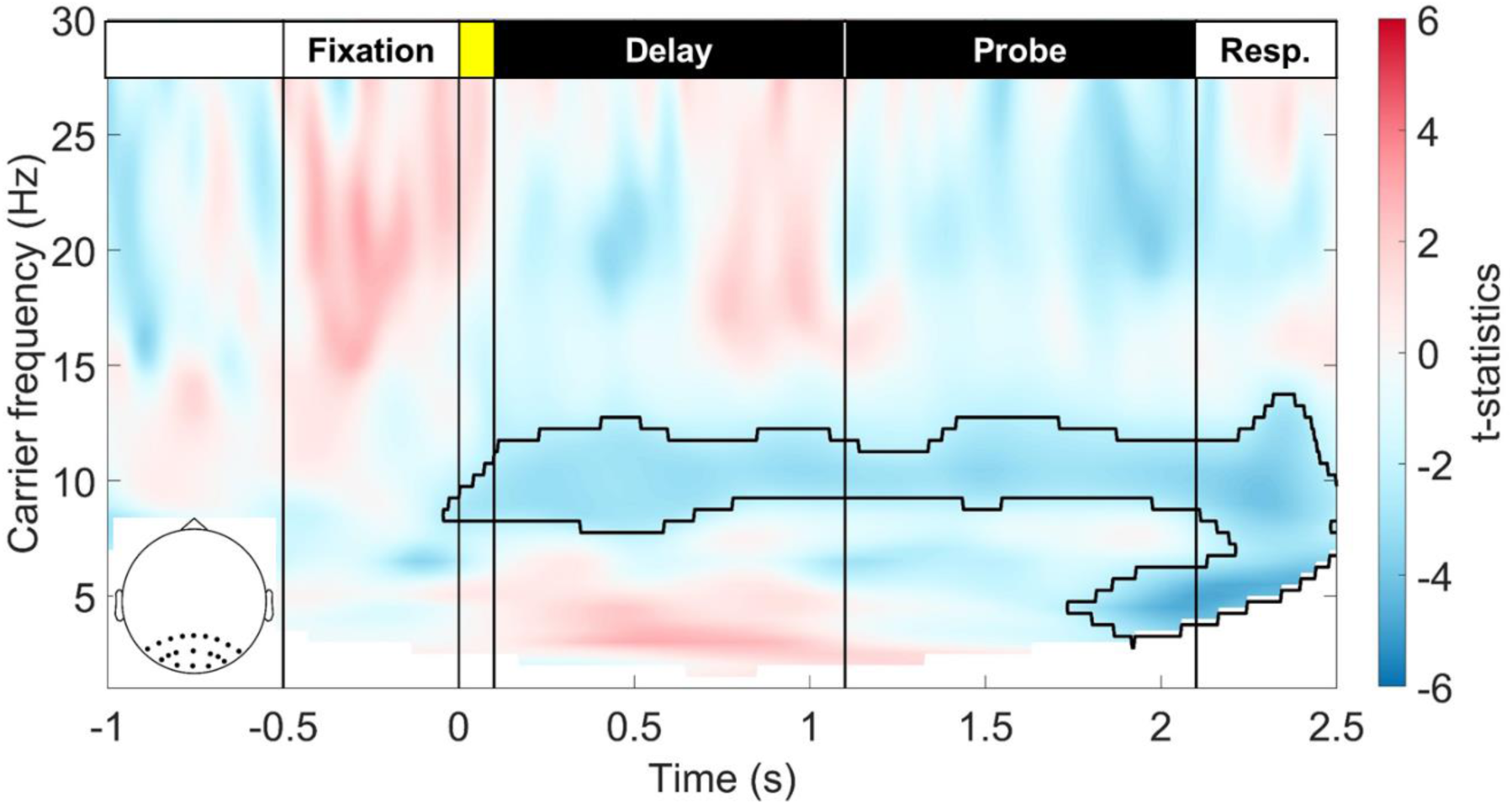
Correlations strengths between WM precision and the parieto-occipital *CF-t* spectrum using the Morlet wavelet transforms. The same GLM analysis was as the parieto-occipital HHS was replicated for the wavelet spectrum, but only the sustained negative alpha cluster was identified.

**Figure S5.**
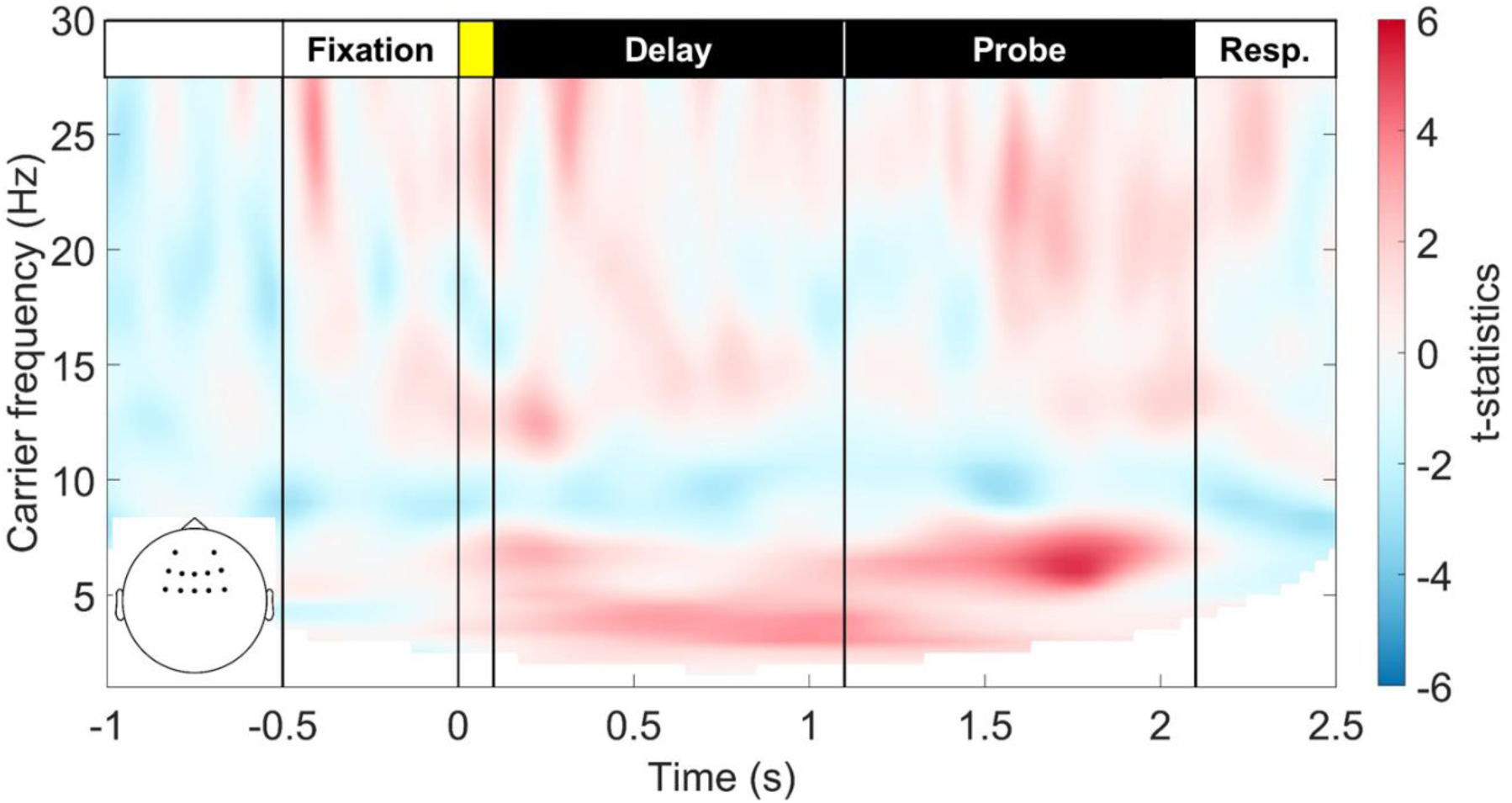
Correlations strengths between WM precision and the frontal *CF-t* spectrum using the Morlet wavelet transforms. No significant cluster was identified using the same GLM analysis as the Fig. 6.

**Figure S6.**
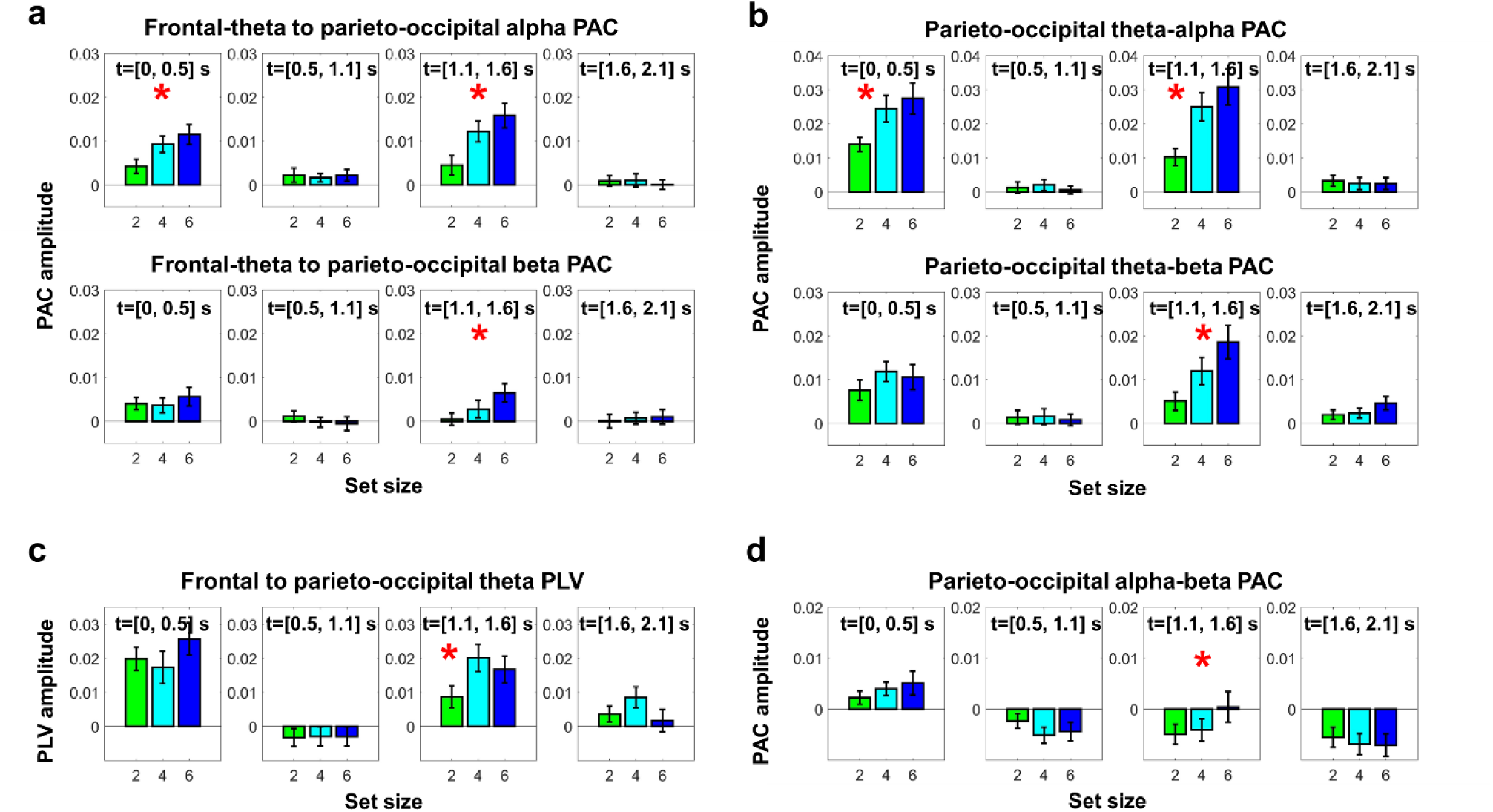
Box plots of relevant local and inter-regional PACs and PLVs for the Figure 7. A repeated-measures ANOVA was applied to each box plot to evaluate the load effect. The set-size 1 condition served as the baseline for comparison. Red asterisks indicated significant load effects after FDR correction. (a) The long-range PAC between frontal theta phase and parieto-occipital alpha/beta amplitudes. (b) The local PAC between parieto-occipital theta phase and alpha/beta amplitudes. (c) the long-range phase synchronization between frontal and parieto-occipital theta oscillations. (d) The local PAC between parieto-occipital alpha phase and beta amplitudes. All significant effects indicated an enhancement from set-size 1 to set-size 6. The only exception is the parieto-occipital alpha-beta PAC, in which the PAC was the smallest in the set-size 2 condition and gradually increased to zero level.

